# RNA G-quadruplexes mark repressive upstream open reading frames in human mRNAs

**DOI:** 10.1101/223073

**Authors:** Pierre Murat, Giovanni Marsico, Barbara Herdy, Avazeh Ghanbarian, Guillem Portella, Shankar Balasubramanian

**Author notes:** Correspondence to: Shankar Balasubramanian.

## Abstract

RNA secondary structures in the 5’ untranslated regions (UTRs) of mRNAs have been characterised as key determinants of translation initiation. However the role of non-canonical secondary structures, such as RNA G-quadruplexes (rG4s), in modulating translation of human mRNAs and the associated mechanisms remain largely unappreciated. Here we use a ribosome profiling strategy to investigate the translational landscape of human mRNAs with structured 5’ untranslated regions (5’-UTR). We found that inefficiently translated mRNAs, containing rG4-forming sequences in their 5’-UTRs, have an accumulation of ribosome footprints in their 5’-UTRs. We show that rG4-forming sequences are determinants of 5’-UTR translation, suggesting that the folding of rG4 structures thwarts the translation of protein coding sequences (CDS) by stimulating the translation of repressive upstream open reading frames (uORFs). To support our model, we demonstrate that depletion of two rG4s-specialised DEAH-box helicases, DHX36 and DHX9, shifts translation towards rG4-containing uORFs reducing the translation of selected transcripts comprising proto-oncogenes, transcription factors and epigenetic regulators. Transcriptome-wide identification of DHX9 binding sites using individual-nucleotide resolution UV crosslinking and immunoprecipitation (iCLIP) demonstrate that translation regulation is mediated through direct physical interaction between the helicase and its rG4 substrate. Our findings unveil a previously unknown role for non-canonical structures in governing 5’-UTR translation and suggest that the interaction of helicases with rG4s could be considered as a target for future therapeutic intervention.

## INTRODUCTION

While all cellular mRNAs use the same translation machinery, the dynamics and efficiency of eukaryotic translation can vary considerably between mRNAs. For example, translation rates in eukaryotes can differ over a 100fold range.^1,2^ This disparity can be explained by the presence of *cis*-regulatory RNA elements modulating the different steps of protein synthesis.^3,4^ While the accessibility of the 5’-cap and the presence of small upstream open reading frames (uORFs) can affect the rate of translation initiation,^5^ codon choice and stop-codon read-through can affect elongation and termination rates respectively.^6^ Despite known examples of regulation during elongation and termination, initiation is the main rate limiting step.^7,8,9^

Canonical eukaryotic translation typically follows a scanning model of initiation, as the m7G cap recruits specific eukaryotic translation initiation factors (eIFs) and the 40S ribosomal subunit to the 5’ end of the mRNA. The assembled 43S preinitiation complex (PIC) then scans the 5’-UTR in the 3’-direction until an initiation codon is encountered. To achieve this, the PIC needs to navigate through regions of the mRNA that contain stable secondary structures and bound proteins. This requires the action of helicases that can either unwind secondary structures or remodel the PIC to facilitate its ability to overcome impediments.^10^ The best-characterised human helicases required for translation initiation comprise the DEAD-box helicases eIF4A^11,12^ (eukaryotic translation initiation factor 4A, also known as DDX2) and DDX3^13^; and the DEAH-box helicases DHX29^14^ and DHX9^15^ (also known as RNA helicase A or RHA). Transcriptome-wide characterisation of human eIF4A^11,12^ and the yeast helicases, eIF4A/B and Ded1,^16,17^ suggests that the eukaryotic translation machinery exploit RNA secondary structures within 5’-UTRs to discriminate between particular mRNA transcripts. Non-canonical secondary structures, such as Hoogsteen-paired G-quadruplexes (rG4s), have been shown to impair translation and were proposed to impede 5’-UTR scanning.^18,19,20^ Nevertheless the nature and extent of human mRNAs that are regulated by rG4s have not been defined. Herein we report transcriptome-wide studies that assess how rG4s affect translation and allow identifying transcripts controlled by two rG4-specific helicases.

## RESULTS

### rG4s within 5’-UTR are *cis*-regulatory elements of translation efficiency

To identify candidate transcripts regulated by secondary structures in their 5’-UTR, we used a transcriptome-wide approach to determine the efficiency of translation of mRNAs in HeLa cells. We adopted a strategy previously used to map ribosome-protected mRNA sequences in cycloheximide (a translation elongation inhibitor)-treated cells (see **Methods**) to define the human translational landscape.^21,22^ Our ribosome-profiling assay generated a 28 nucleotide (nt) ribosome-protected fragments (RPFs) peak (**Extended** Figure 1a). In parallel, we performed matched transcriptional mRNA sequencing (mRNA-seq) and defined the translation efficiency (TE) for each transcript by normalising the ribosome footprint frequency to total transcript abundance (transcript per millions (TPM) using signal over CDSs). Across replicates (n = 2), TE values from our ribosome profiling analysis were highly reproducible (Spearman correlation = 0.935, **Extended** Figure 1b). While RPF counts span 5 orders of magnitude, mRNA counts span 4 orders of magnitude (**Extended** Figure 1c) revealing a wide dynamic range in genome-wide translational efficiencies. The transcriptome-wide distribution of TE (Figure 1a) showed a skewed distribution (skewness = −1.069) towards inefficiently translated transcripts. Interestingly, genes that were efficiently translated (4^th^ quartile of log_2_ TE distribution) are associated with the maintenance of basic cellular functions, while genes that were inefficiently translated (1^st^ quartile) are associated with cancer pathways (Figure 1b). This observation suggests specific mechanisms in HeLa cells that maintain the TE of cancer genes at a level that does not exceed a threshold necessary for cellular homeostasis.^23^

**Figure 1.**
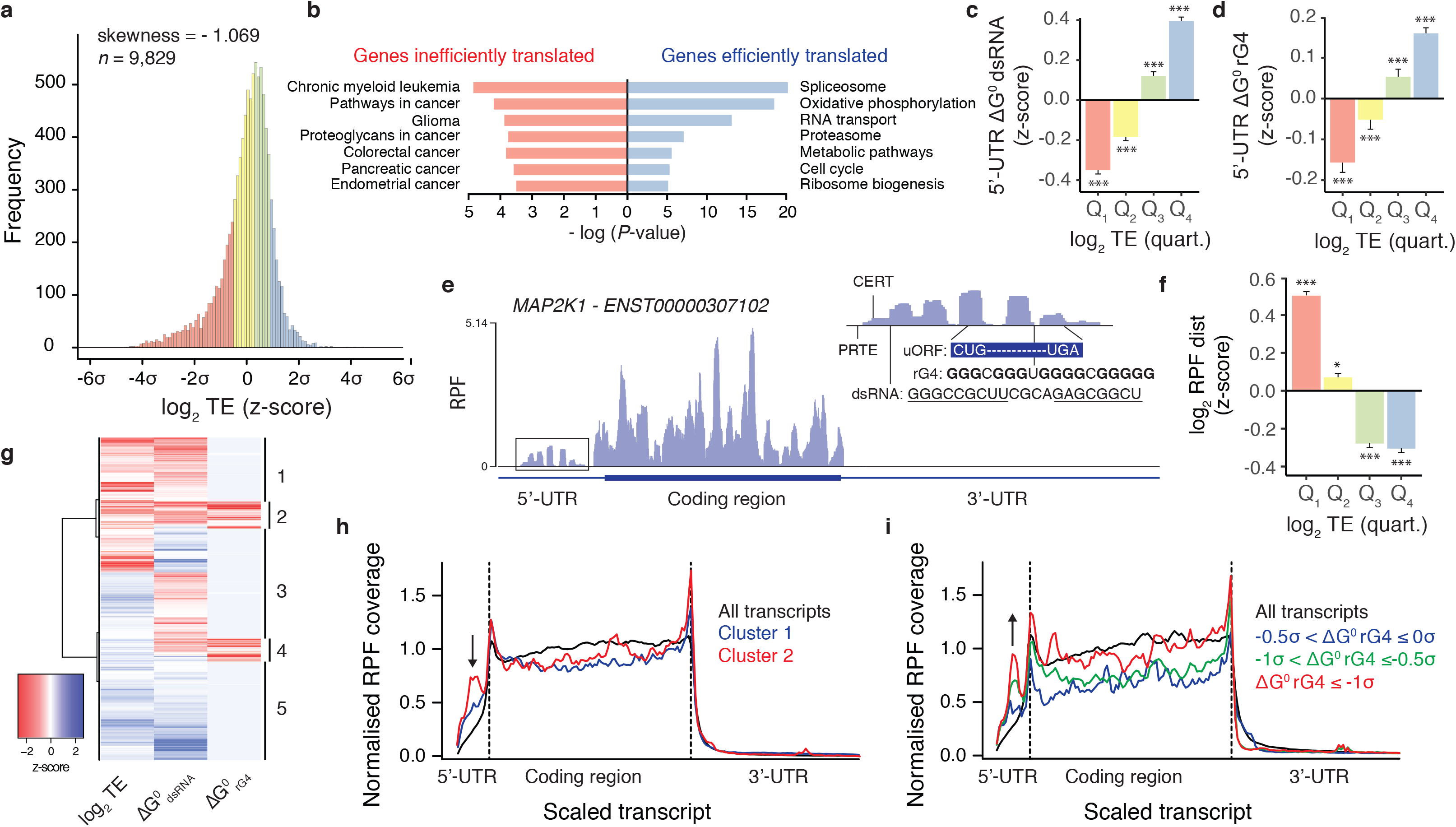
I RNA secondary structures within 5’-UTR alters ribosome distribution and impede translation of associated CDSs. **a)** The transcriptome-wide distribution of translation efficiency of human mRNAs shows a skew towards inefficiently translated transcripts. **b)** Pathway analysis of inefficiently (1^st^ quartile of TE distribution, red) and efficiently (4^th^ quartile, blue) translated genes. **c)** dsRNA and **d)** rG4 associated predicted folding energies of 5’-UTRs of human transcripts binned according to their TE (1^st^ to 4^th^ quartile of TE distribution). Folding energies are expressed as z-scores of the minimum free energies normalised by the length of the 5’-UTRs. **e)** Ribosome density within *MAP2K1* mRNA showing high ribosome occupancy in the 5’-UTR. As shown in the insert, RPFs in 5’-UTR colocalize with potential *cis*-regulatory elements of translation initiation. Excess of RPF in 5’-UTR correlates with inefficient translation of associated CDSs: **f)** RPFdist, expressed as z-score, of human mRNAs binned according to their TE (1^st^ to 4^th^ quartile of TE distribution). **g)** Hierarchical clustering analysis of human transcripts according to their TE and length-normalised dsRNA and rG4 predicted folding energies of 5’-UTRs. The heatmap report the z-scores of the three variables. **h)** Ribosome distribution for transcripts of cluster 1 (blue) and cluster 2 (red), *i.e*. transcripts with low TE and associated with predicted stable dsRNA or rG4 structures respectively. **i)** Ribosome distribution for transcripts of cluster 2 when binned for increasing predicted rG4 stability, – 0.5σ < ΔG^0^rG4 ≤ 0σ (blue line), −1σ < ΔG^0^rG4 ≤ −0.5σ (green line) and ΔG^0^rG4 ≤ −1σ (red line), where ΔG^0^rG4 is the z-score of 5’-UTR length-normalised rG4 predicted minimum free energy. Ribosome footprint coverage and transcript length are normalized; dotted lines indicates annotated translation start and stop sites and the arrows highlight the presence of RPFs in 5’-UTR. Data reported in (**c**), (**d**) and (**f**) are means ± s.e.m, *P*-values were assessed using one-tailed Mann-Whitney nonparametric tests and represent statistical difference between the binned population and the rest of the population. \**P* < 0.05, \*\*\**P* < 0.001.

TE correlated with the length, GC-content and Gscore, a measure of guanine richness and skewness,^24^ of 5’-UTRs (see **Supplementary Information** for a complete definition of the mRNA features discussed in this manuscript, **Extended** Figure 1d-f) indicating that translational control of gene expression is to a great degree encoded in 5’-UTR sequences.^1^ As expected, the 5’-UTR of inefficiently translated transcripts were enriched in known *cis*-regulatory elements, such as Cytosine Enriched Regulator of Translation (CERT),^25^ Pyrimidine-rich translation element (PRTE)^26^ or uORFs^27^ (**Extended** Figure 1g-j). Secondary structures in 5’-UTRs also played an important role in modulating TE, as exemplified by the enrichment of rG4-forming sequences in inefficiently translated transcripts or the negative impact of canonical double-stranded RNA (dsRNA) structures closed to the 5’-cap (**Extended** Figure 1k-l). When normalised for length, reduced predicted dsRNA and rG4 secondary structures folding energy of 5’-UTRs, *i.e*. more structured 5’-UTRs, correlated with inefficient translation (Figure 1c-d). Taken together, these observations demonstrate that the presence of known *cis*-regulatory elements correlates with diminished TE on a transcriptome-wide scale. It is noteworthy that we observe a general trend linking predicted stable rG4s in 5’-UTR with low TEs, an effect that was previously reported only for a limited number of transcripts.^20^

Manual inspection of inefficiently translated mRNAs revealed a ribosome distribution shifted to a higher density within 5’-UTRs. For instance, Mitogen-Activated Protein Kinase Kinase 1 (*MAP2K1*, also known as *MEK1*), an essential component of the ERK signal transduction pathway and frequently mutated in cancer,^28^ exhibited high ribosome occupancy within the 5’-UTR (Figure 1e). Ribosomes in *MAP2K1* 5’-UTR co-localised with *cis*-regulatory elements of translation suggesting an impact of ribosome distribution and 5’-UTR translation on TE. When considering RPFdist, a proxy of 5’-UTR translation,^29^ defined by the ratio of RPF reads within the 5’-UTR relative to reads within the annotated downstream CDS (see Methods), we found that an excess of RPF in 5’-UTR, *i.e*. high RPF value, is associated with low translation efficiency (Figure 1f). To support this point, when considering transcripts with higher or lower than average RPFdist values, we found *Pearson* correlations between RPFdist and TE of −0.344 and −0.070 respectively (**Extended** Figure 1m). We found that transcripts marked by high ribosomal coverage along the 5’-UTR (4^th^ quartile of log_2_ RPFdist, **Extended** Figure 2a) are characterised by low GC content but high local skew in guanine versus cytosine composition (**Extended** Figure 2b-c). We also found an enrichment of potential G-quadruplex forming sequences and reduced folding energies of the predicted secondary structures in this population of transcripts (**Extended** Figure 2d-f). To decipher the contribution of rG4 from canonical dsRNA structures, we devised a hierarchical clustering approach (see Methods) to identify groups of transcripts of similar TE and 5’-UTR folding energies. This approach allowed two clusters to be identified, clusters 1 and 2 (Figure 1g), associated with low TE and stable predicted secondary structures in their 5’-UTR. While cluster 1 was characterised by stable 5’-UTR dsRNA structures, cluster 2 was enriched in predicted stable rG4 structures. It is noteworthy that the presence of rG4 and dsRNA tends to be mutually exclusive (**Extended** Figure 2g). Transcripts of clusters 1 and 2 showed altered ribosome footprint distribution as compared to the global transcript population (Figure 1h and **Extended** Figure 2h) with RPF coverage increased in 5’-UTRs and decreased in CDSs. It is noteworthy that the presence of rG4 has greater impact on RPF distribution than dsRNA structures (Figure 1h). Notably we found that RPF density along 5’-UTR increased when considering rG4 structures of increasing predicted stability (Figure 1i) suggesting a direct role for rG4s in the translation of 5’-UTRs.

### rG4-forming sequences are determinants of 5’-UTR translation

Although the presence of predicted rG4 structures correlated with increased RPF density within 5’-UTR and reduced TE, rG4s do not fully account for the observed variation in our dataset (as exemplified by the correlation between the presence of CERT and PRTE elements and RPFdist, see **Extended** Figure 2i-j). We thus sought to devise a quantitative model that integrates mRNA features, such as mRNA abundance, 5’-UTR secondary structures, 5’-UTR and CDS length, sequence composition statistics, the presence of AUG and non-AUG uORFs and known *cis*-regulatory elements, with the goal of predicting RPFdist measurement, *i.e*. 5’-UTR ribosome occupancy. A principal component analysis (see **Methods** and **Supplementary Information**) based on a subset of these features showed that rG4-containing transcripts defined a distinct group of transcripts (Figure 2a). It is noteworthy, that dsRNA and rG4 features are separated by the second component suggesting that they contribute to ribosome distribution independently (**Extended** Figure 3a-b). The subset of transcripts characterised by discrete rG4 predicted structures marking their 5’-UTR (defined on the PCA by Dim.1 ≥ 0 and Dim.2 ≤ 0) showed an increased ribosome density within their 5’-UTRs compared to the rest of the population (Figure 2b). We then constructed regression models (see **Supplementary Information**) to predict the RPFdist values of two sets of transcripts based on a list of potential predictors. The first set of transcripts included the 8,024 transcripts expressed in HeLa cells with both 5’-UTRs and 3’-UTRs annotated, while the second group was the subset of 1,841 transcripts displaying clear signature associated with rG4 structures within their 5’-UTRs (defined by the PCA with Dim.1 ≥ 0 and Dim.2 ≤ 0). Our model trained on the global population of transcripts explained ~56% of the variance in RPFdist (**Extended** Figure 3c), whereas our model trained on the rG4-containing 5’-UTR subset explained ~65% of the variance in RPFdist (Figure 2c). To gauge the overall performance of our models, we challenged them against three independent test sets and found consistent prediction abilities (**Extended** Figure 3d-k). This result shows that considering predicted rG4 structures substantially improve the prediction of ribosome distribution. This point is supported by the fact that a model selected using only rG4-based predictors accounted for 14.4 ± 3.4 % and 32.1 ± 8.4 % (mean ± s.d. over 10 resampling steps) of the RPFdist variance of the global population of transcripts and the rG4-containing 5’-UTR subset respectively (Figure 2d, Extended Figure 3l). Moreover, our model showed that rG4-based predictors explained RPFdist variance as well as uORF-based predictors (32.0 ± 7.4 %, mean ± s.d. over 10 resampling steps, Figure 2d). Our findings demonstrate that rG4 structures are determinants of ribosome distribution.

**Figure 2.**
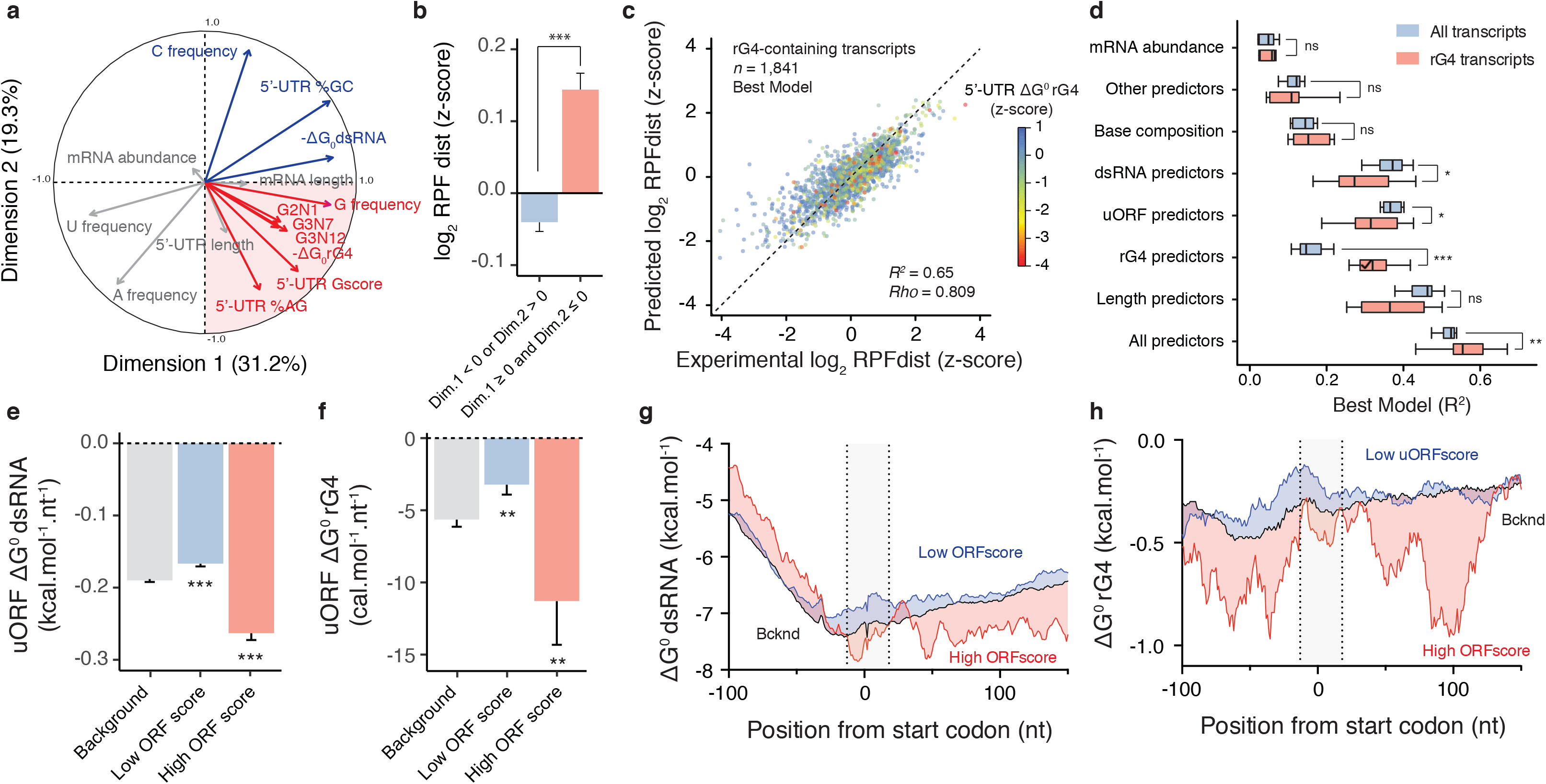
I RNA G-quadruplexes are determinants of 5’-UTR translation. **a)** Principal component analysis of human transcripts using features describing mRNA abundance, 5’-UTR secondary structures, 5’-UTR and mRNA length and 5’-UTR sequence composition statistics. The first two principal components, explaining ~50% of the variance, separate features describing rG4 structures (red quadrant) from features describing dsRNA structures (see also **Extended** Figure 3a). **b)** The subset of transcripts displaying clear signature of rG4 structures within their 5’-UTRs (defined by the PCA with Dim.1 ≥ 0 and Dim.2 ≤ 0) are characterised by an increased ribosome occupancy in 5’-UTRs when compared to the rest of the population. **c)** A statistical model with as few as 32 predictors explains 65% of the RPFdist variation observed in the rG4-containing subset of transcripts. **d)** Performance of models selected using a subset of predictors on either all transcripts or the rG4-containing subset of transcripts. A model selected using rG4-based predictors only can account for 32.1 ± 8.4 % of the observed RPFdist variance in the rG4-containing subset of transcripts making rG4-based predictors as informative as uORF-based predictors (32.0 ± 7.4 %, mean ± s.d. over 10 resampling steps). Reduced predicted (**e)** dsRNA and (**f)** rG4 secondary structures folding energies are associated with translated uORF, *i.e*. high ORFscore uORFs (ORFscore ≥ 6). Folding energies are expressed as the minimum free energies normalised by the length of the uORFs. Secondary structure potential around the start codons of translated uORFs. **g)** dsRNA and (**h)** rG4 predicted folding energies were calculated per position using a sliding window of 35 nt. The grey area represents the size of a 80S ribosome on an upstream start codon. Data in (**b**), (**e**) and (**f**) are means ± s.e.m, *P*-values were assessed using one-tailed Mann-Whitney nonparametric tests and compare the reported condition to background. Central black lines in (**d**) represent the medians and the other black lines represent quartile boundaries. *P*-values were assessed using Kolmogorov–Smirnov nonparametric tests. ns: non significant, \**P* < 0.05, \*\**P* < 0.01, ***P < 0.001.

In order to assess whether ribosomes within rG4-containing 5’-UTRs are translating or stalled/poised, we sought to identify signature of active translation within 5’-UTRs. We therefore assessed the periodicity of ribosome movement on 5’-UTRs to identify actively translated uORFs (see **Methods**). Given a putative uORF (5’-leader sequence beginning with either AUG, CUG, UUG and GUG start codons in frame with a stop codon within the 5’-UTR), we quantified the number of RPFs in each frame and calculating an ORFscore^30^ reflecting whether RPFs were uniformly distributed or preferentially accumulated in the first frame. Among all potential uORFs, we identified 7,650 uORFs displaying enough coverage to robustly assess an ORFscore. Among these, we identified 1,522 uORFs with a low score (0 ≤ ORFscore < 6) and 274 uORFs with a high score (ORFscore ≥ 6). Low ORFscores are associated with uORFs that are unlikely to be translated (low consistency between RPF distribution and the frame of the uORF), while high ORFscores are associated with periodic and in frame translation of the uORF. We observed that translated uORFs (High ORFscore) were on average longer than all potential uORFs (**Extended** Figure 4a).^31^ More interestingly, we found that uORFs with a high ORFscore are characterised by high GC content and G-score (**Extended** Figure 3b-c) and are enriched in known *cis*-regulatory elements of translation, such as CERT motifs or rG4 motifs (**Extended** Figure 4d-g). When normalised for length, reduced predicted secondary structure folding energies of uORFs also correlated with active 5’-UTR translation (Figure 2e-f). These observations support the role of RNA secondary structures in stimulating 5’-UTR translation. We then examined the initiation context of translated uORF starts to assess how secondary structures contribute to the translation of uORFs. While there was no sign of particular GC richness in the vicinity of upstream start codons of translated uORFs, higher G-scores were observed in the vicinity of the start codon of high ORFscore uORFs when compared to low ORFscore uORFs (**Extended** Figure 4j-k). This observation was mirrored by the enrichment of predicted rG4 structures, but not of dsRNA structures, directly before and after the start codons of high ORFscore uORFs (Figure 2g-h). These observations suggest that rG4 structures promote 5’-UTR translation probably by arresting or slowing PIC scanning in the vicinity of upstream start codons, hence promoting 80S ribosomes formation.

### DHX9 and DHX36 are associated with polysomes and bind RNA G-quadruplexes

To demonstrate that rG4 structural motifs, rather than the intrinsic guanine-richness of the sequences, affect both ribosome distribution and translation efficiency, we sought to identify protein factors associated with rG4s that modulate the effect of predicted rG4 structures. We envisaged that such factors would be associated with polysomes, *i.e*. actively translated mRNAs, and bind to the rG4 structural motif. Therefore, we performed polysome profiling coupled with mass spectrometry (see **Methods**) to identify suitable candidates. HeLa cell lysates were loaded on sucrose density gradients and fractions corresponding to supernatant, monosomes (40S, 60S and single 80S) and polysomes were recovered by fractioning the ultracentrifugated gradients (**Extended** Figure 5a). Isolated fractions were resolved by gel electrophoresis and high-molecular weight protein complexes were analysed by mass spectrometry. This approach allowed identifying proteins associated with polysomes and probing their repartition within the components of the lysate (**Extended** Figure 5b). Using a functional analysis, we found a significant enrichment of DEAD- and DExH-box helicases in the polysomes fractions (**Extended** Figure 5c). A quantitative estimate (Figure 3a) revealed that three out of the six paralogs of the DEAH-box/RHA family^32^ of helicase were enriched in the polysomal fractions, namely DHX9, DHX30 and DHX36, suggesting a strong link between this helicase family and translation regulation. We then assessed the enrichment of each helicase in the different fractions of the polysome gradient by immunoblotting (Figure 3b). We confirmed that DHX9, DHX30 and DHX36 were present in both the heavy and light polysome fractions while DHX29 and DHX57 were enriched in the monosome fractions. The two other helicases from the family, TDRD9 and YTHDC2, were not found associated with either mono or polysomes. We then interrogated the ability of these helicases to bind RNA structures by performing affinity enrichments (see **Methods**). We found an rG4 probe enriched DHX9, DHX36 and DHX57, while a similar probe mutated to abrogate rG4 formation showed now enrichment. A probe folding into a stem loop structure was not able to enrich for the helicases (Figure 3b). Taken together, these results suggest that DHX9 and DHX36 are appropriate candidates to study the role of rG4 on translation since both helicases are present in rG4 / ribonucleoprotein complexes and are associated with translated mRNAs. This conclusion is further supported by the ability of DHX9 and DHX36 to unwind rG4s *in vitro* and in modulating the translation of selected structured mRNAs.^33,15,34,35^

**Figure 3.**
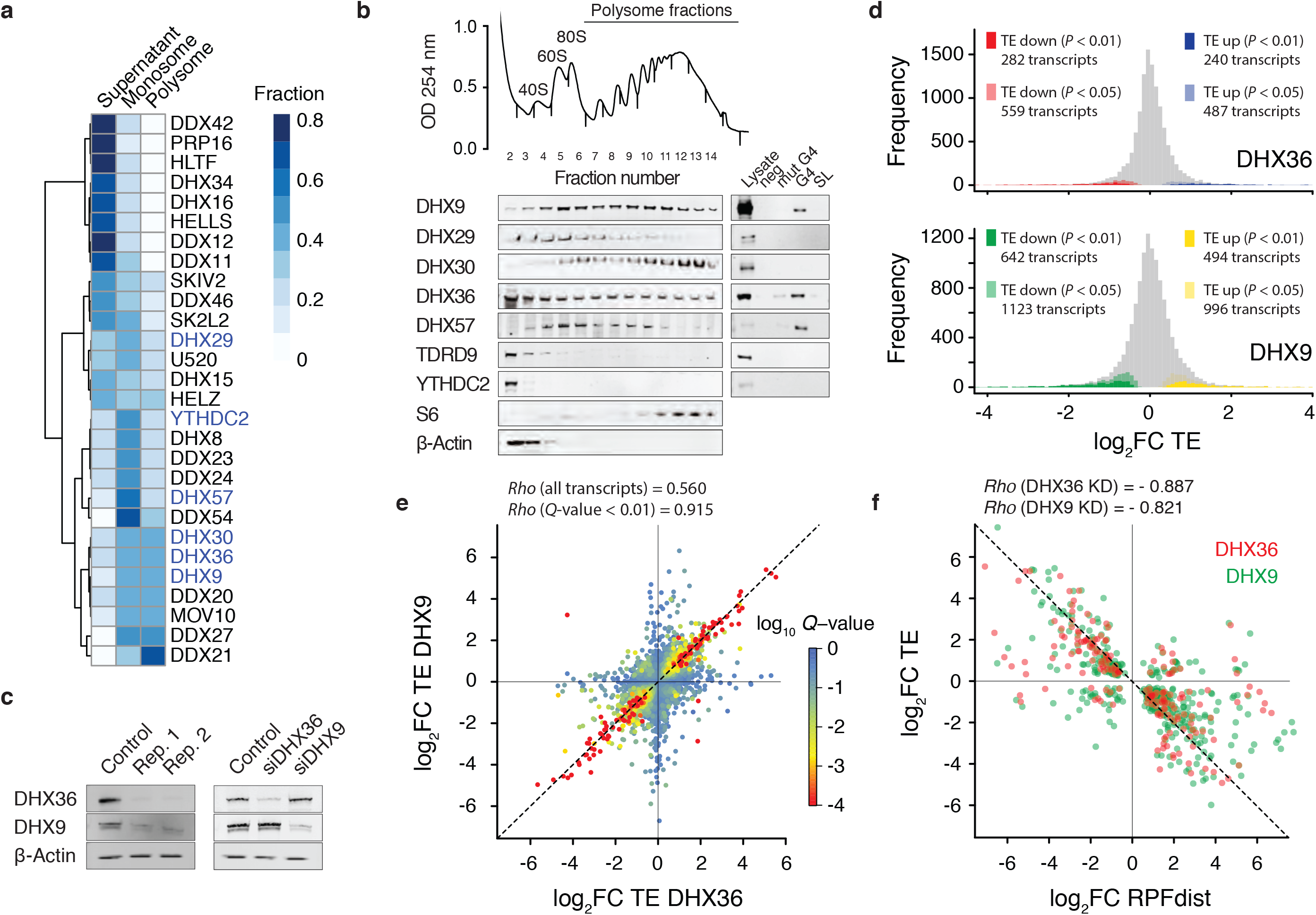
I Polysome and ribosome profiling defines the role of DHX36 and DHX9 on translation. **a)** Polysome profiling of HeLa cytoplasmic extract coupled with mass spectrometry allowed estimating the enrichment of different helicases in monosome or polysome fractions. Duplicates were used to quantitatively estimate the presence of a given helicase in each fraction. A hierarchical cluster analysis reporting the distribution of human helicases within mono/polysomes is shown. **b)** Immunoblots of a polysome profile probing for members of the DEAH-box/RHA helicase family (left blot) confirmed the presence of DHX36 and DHX9 in both light and heavy polysomes. On the right is reported the result of affinity purifications using indicated biotinylated nucleic acids probes showing the presence of DHX9, DHX36 and DHX57 in rG4 / ribonucleoprotein complexes. Lysate: 30ug of total protein, Neg: empty beads, mut G4: mutated rG4-forming sequences, rG4: rG4-forming sequences, SL: stem-loop forming sequence. **c)** Immunoblots of siRNA-treated cells probing DHX36 and DHX9 show that siRNA depletion of both helicases is reproducible and selective. **d)** Ribosome profiling allowed assessing change in TE upon helicase depletion. Are reported the frequency distributions of the ratio of TE in DHX36 (top) and DHX9 (bottom) depleted cells over TE in control (*n* = 3 replicates). **e)** Comparison of fold changes of TE upon DHX36 and DHX9 depletion. *Q*-values were calculated by combining *P*-values using Fisher’s method. **f)** Anticorrelation between fold changes of TE and fold changes of RPFdist upon depletion of DHX36 (red) or DHX9 (green) for transcripts showing significant change in TE (*Q*-value < 0.05).

### Translational landscapes of DHX36- and DHX9-deficient cells

To understand how both helicases contribute to the translation of rG4-containing mRNAs, we performed ribosome profiling of HeLa cells depleted in DHX9 and DHX36 by siRNA. Knockdowns of both helicases (see **Methods**) were found efficient and selective (Figure 3c). We assessed the TE of all HeLa transcripts, in triplicates for both knock down conditions and a control experiment (pool of non-targeting siRNAs). To evaluate the TE change between the DHX36, DHX9 and control samples, we then calculated the ratios TE_DHX36_ / TE_control_ and **TE_DHX9_** / TE_control_. Changes in TE were highly reproducible across triplicates (**Extended** Figure 6a-b). We then contrasted genome-wide transcriptional and translational differences (**Extended** Figure 6c-f), and found that change in TE upon helicase depletion correlated with change in RPF signal (Spearman correlation = 0.762 and 0.744 for DHX36 and DHX9 samples respectively) rather than change in total RNA signal (Spearman correlation = −0.239 and −0.299 for DHX36 and DHX9 samples respectively), indicating a minimal impact of transcriptional variation in our measure of TE variation. We identified 1,026 and 2,119 transcripts (using a cut-off at *P* < 0.05; see **Methods**) that were significantly affected by the depletion of DHX36 and DXH9 respectively (Figure 3d). Interestingly, change in TE upon depletion of DHX36 strongly correlated with change in TE upon depletion of DHX9 (Pearson correlation = 0.915 for mRNAs both affected by both DHX36 and DHX9 depletion, Figure 3e). We also found a strong anticorrelation between change in TE and in RPFdist (Figure 3f, Pearson correlation = −0.887 and −0.821 for DHX36 and DHX9 depletion respectively), showing that change in TE is accompanied with a significant shift in RPF distribution. Again, changes in RPFdist in both experiments were strongly correlated (Pearson correlation = 0.915, Extended Figure 6g). Overall, these observations suggest that both helicases may share a common mechanism to regulate translation

### G-quadruplexes mark DHX36- and DHX9-dependent mRNAs

Due to the high overlap between the mRNAs whose translation efficiency was affected (**Extended** Figure 6h) and the high overlap between the mRNAs whose patterns of ribosomal occupancies were affected by the depletion of both helicases (**Extended** Figure 6i), we defined two groups of mRNAs: the first group, hereafter named TE_down_ – RPFdist_up_, is characterised by diminished TE and increased RPFdist, while the second group, hereafter named TE_up_ – RPFdist_down_, is characterised by increased TE and diminished RPFdist upon the depletion of both helicases (see **Methods**). The TE_down_ – RPFdistup group included 282 transcripts, the TE_up_ – RPFdist_down_ group includes 195 transcripts, while the background list (whose TE is not affected by the depletion of the helicases) included 946 transcripts.

We compared the TE_down_ – RPFdist_up_, TE_up_ – RPFdist_down_ and background groups and found that the 5’-UTR of both transcript groups affected by the depletion of the helicases were longer than the 5’-UTR of the background transcripts (**Extended** Figure 6j). When normalised for 5’-UTR length, a reduced 5’-UTR predicted rG4 folding energy was found in the TE_down_ – RPFdist_up_ but not in the TE_up_ – RPFdist_down_ group (**Figure 4a**). We also observed an enrichment of rG4s in the 5’-UTR of the TE_down_ – RPFdist_up_ group (**Extended** Figure 6k). Interestingly, no differences in predicted dsRNA folding energies were observed (Figure 4b). These findings suggest that 5’-UTR length and rG4 secondary structures make significant contribution to changes in TE upon depletion of the helicases.

**Figure 4.**
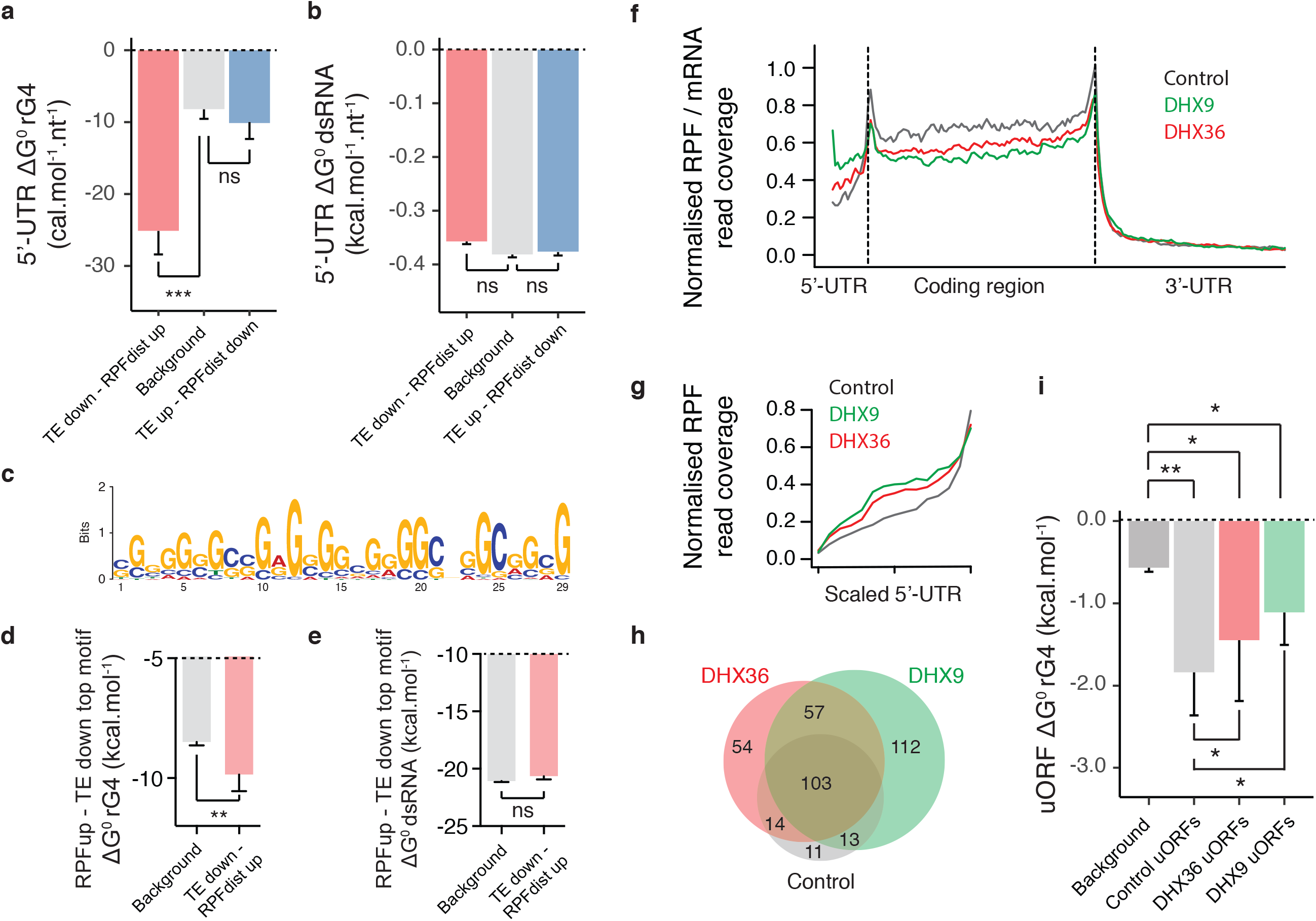
I Translation is shifted towards rG4-containing 5’-UTRs in DHX36 and DHX9 depleted cells. rG4s mark the 5’-UTR of DHX36-and DHX9-dependent mRNAs. Are reported the 5’-UTR length-normalised (**a**) rG4 and (**b**) dsRNA secondary structures predicted folding energies of the TE_down_ – RPFdist_up_ and TE_up_ – RPFdispdown groups compared to background transcripts (see main text for description of the different sets of transcripts). **c)** Most enriched motif in the 5’-UTR of the TE_down_ – RPFdist_up_ group (*P* < 2.2 × 10^−16^, Fisher exact test). The same motif was found depleted in the 5’-UTR of TE_up_ – RPFdisp_down_ group (*P* = 2.4 × 10^−2^, Fisher exact test). **d)** rG4 and (**e**) dsRNA secondary structures predicted folding energies of the TE_down_ – RPFdist_up_ top motif in the background or TE_down_ – RPFdist_up_ transcripts. Sequences corresponding to the identified motif ± 10 nt were considered to reflect the influence of 5’-UTR base composition on local predicted structures. **f)** Ribosome distribution normalised by mRNA signal, describing local translation efficiency, of the TE_down_ group (380 transcripts with combined *Q*-value ≤ 0.05) and (**g**) ribosome distribution of the RPFdist_up_ group (548 transcripts with combined *Q*-value ≤ 0.05) in control (black), DHX36 (red) and DHX9 (green) depleted cells. The plots show increase and decrease of translation of 5’-UTRs and associated CDSs respectively. Ribosome footprint, mRNA signal coverages and transcript length are normalized; dotted lines indicates annotated translation start and stop sites. **h)** Venn diagram depicting total number of detected high ORFscore uORFs in control (black), DHX36 (red) and DHX9 (green) depleted cells. Depletion of the helicases induces the translation of new uORFs. i) Predicted rG4 structure folding energies of detected high ORFscore uORFs in control (grey) or DHX36 (red) and DHX9 (green) depleted cells. The background set (black) represents uORFs with negative ORFscore in control cells. Data are means ± s.e.m, *P*-values were assessed using one-tailed Mann-Whitney nonparametric tests. ns: non significant, \**P* < 0.05, \*\**P* < 0.01, ***P < 0.001.

Using the MEME algorithm,^36^ we looked for enrichment of motifs in each group (**Extended Figure 7**). While the TE_down_ – RPFdist_up_ group was characterised by the enrichment of short (12 to 30nt) GC-rich motifs, the TE_up_ – RPFdist_down_ group was enriched in AT-rich motifs (**Extended Figure 7a-b**). The motifs enriched in the TE_down_ – RPFdist_up_ group were depleted in the TE_up_ – RPFdist_down_ group (**Extended Figure 7c-d**). The TE_down_ – RPFdist_up_ top motif (Figure 4c) showed a skew in guanine composition and was found to overlap with predicted G-quadruplex forming sequences (**Extended **Figure 7e****). Interestingly, when comparing the context of the sequences matching this motif in the TE_down_ – RPFdist_up_ or the background group, we found that the sequences in the 5’-UTR of the TE_down_ – RPFdist_up_ group were characterised by a skew in purine, and more particularly in guanine, with no bias in GC content (**Extended Figure 7f-h**). This sequence context favours the formation of rG4 structures, which is confirmed by smaller values of rG4 folding energies but similar values of dsRNA folding energies (Figure 4d-e). Taken together these results demonstrate that rG4 structures in 5’-UTRs is a significant determinant of diminished TE caused by impairing DHX36 and DHX9 function.

### Depletion of DHX36 and DHX9 induces shifts in translation

Diminished TE co-occurred with an increase in 5’-UTRs translation as seen on Figure 4f. Ribosome density within 5’-UTRs was respectively increased within the 5’-UTR of the RPFdist_up_ group (Figure 4g). These observations suggests that DHX36 and DHX9 depletion induced a shift of translation towards rG4-containing 5’-UTRs. To confirm this point, we sought to identify uORFs that are activated upon depletion of the helicases. We assigned an ORFscore to each potential uORFs and selected the ones with an ORFscore ≥ 6, as uORFs with RPF patterns consistent with active translation. We detected 223 additional translated uORFs in cells depleted in DHX36 or DHX9 as compared to the control cells (141 translated uORFs, Figure 4h). The detected DHX36- and DHX9-dependent uORFs significantly overlapped (with 57 out of 223 new uORFs overlapping, *P* < 0.01, Fisher exact test) confirming a shared mechanism for controlling 5’-UTR translation. The DHX36- and DXH9-dependent uORFs were characterised by more stable predicted rG4s than uORFs from background, *i.e*. uORF with negative ORFscore in control sample, but less stable than the DHX36- and DHX9-independent uORFs (Figure 4i). This latter result could not be explained by differences in uORF length (**Extended Figure 8a**) and was not observed when considering dsRNA structures (**Extended Figure 8b**). These observations show that metastable rG4s mark DHX36- and DHX9-dependent uORFs and their folding, favoured by the absence of the helicases, shifts translation to 5’-UTRs.

### DHX9 binds RNA-quadruplexes in human cells

To assess whether the helicases modulate TE through direct interaction with rG4 motifs, we used a transcriptome-wide approach to determine the DHX9 RNA-binding targets in HeLa cells. We adapted the Individual-nucleotide resolution UV crosslinking and immunoprecipitation (iCLIP) method^37^ (see **Methods**) to map DHX9 binding sites. The nature of the immunoprecipitated RNA-DHX9 complex was confirmed using control experiments performed under identical conditions but either excluding the UV cross-linking step (Figure 5a) or omitting the DHX9 specific antibody during the immunoprecipitation step (**Extended Figure 9a**). Deep sequencing of immunoprecipitated RNAs allowed the identification of 2,667 transcripts (encoding 2,411 individual genes) bound by DHX9 and the two perfomed replicates showed good reproducibility (**Extended Figure 9b**, *Spearman* correlation = 0.88 between number of reads per transcripts in each duplicate). We identified 5,152 peaks with ~ 2 peaks in average per transcripts. Among these peaks, 7.6 % and 89.3 % were found in 5’-UTRs and 3’-UTRs respectively. The called peaks showed a high overlap between replicates (**Extended Figure 9c**, 97 % of the peaks from the second replicates overlapped with the peaks identified in the first experiment) supporting the robustness of our iCLIP protocol. After multimodal peak splitting, we identified 11,235 individual binding events characterised by discrete peaks of median width of 82 nt (**Extended Figure 9d**).

**Figure 5.**
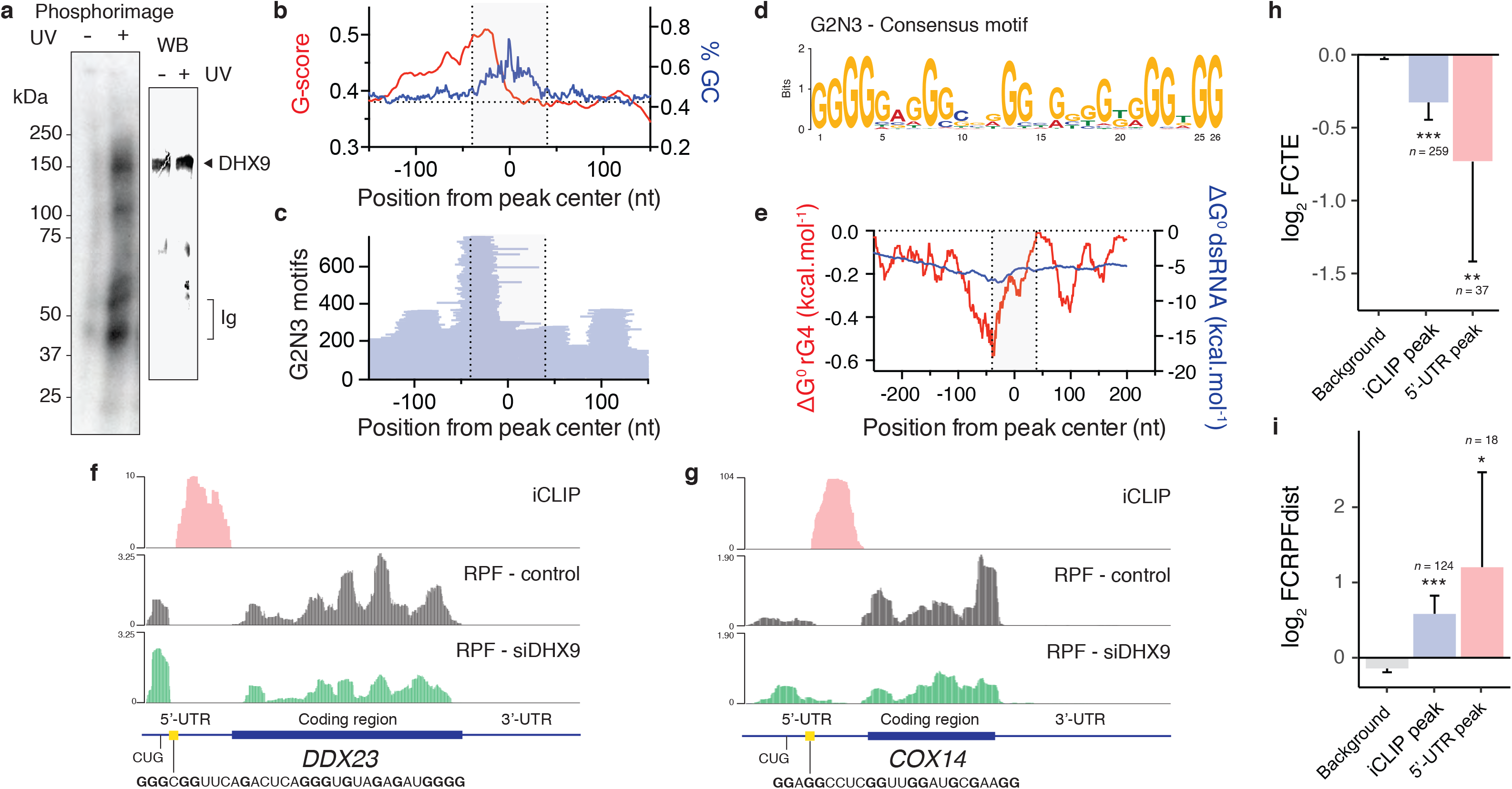
I DHX9 binds downstream rG4 motifs in human cells. **a)** Phosphorimage of SDS gel (left gel) resolving ^32^P-labelled RNAs crosslinked to DHX9. Immunoprecipitated samples prepared from HeLa cell lysates without UV light (254 nm) treatment are shown as a negative control. Immunoblot of the same membrane (right) probing DHX9 confirm the presence of DHX9 in the RNA-protein complex. **b)** Gscore (red) and GC content (blue) of sequences around the centre of iCLIP peaks. **c)** Distribution of rG4 motifs, of the form G2N3 (see text for details), around the centre of iCLIP peaks. The grey area represents the median width of iCLIP peaks. **d)** Consensus motif of rG4-forming sequences found upstream DHX9 iCLIP peaks. **e)** rG4 and dsRNA predicted folding energies within 200 bases from the centre of 5’-UTR DHX9 iCLIP peaks. Minimum folding energies were calculated using a sliding window of 35 nt. Mapped DHX9 binding sites (red) and normalised ribosome density in control (black) and DHX9 depleted cells (green) within the (**f**) *DDX23* and (**g**) *COX14* transcripts. Depletion of DHX9 leads to an increase and a decrease of RPF within the 5’-UTRs, upstream of DHX9 binding sites, and associated CDSs respectively. Fold changes in (**h**) TE and (**i**) RPFdist of transcripts directly bound by DHX9 anywhere (blue) or within 5’-UTR (red). Data are means ± s.e.m, *P*-values were assessed using two-sample Kolmogorov-Smirnov tests and compare the reported condition to background. \**P* < 0.05, \*\**P* < 0.01, \*\*\**P* < 0.001.

We found an enrichement in G and C, and a depletion in A and U residues within these DHX9 binding peaks (**Extended Figure 10a**). These observations suggest that DHX9 binds structured RNA sequences in a cellular context. Interestingly a higher than baseline Gscore was observed upstream of the GC-rich DHX9 binding sites (Figure 5b) suggesting a role for rG4 motifs in DHX9 binding. We then explored whether the relative positioning and frequency of rG4s created a specific signature at DHX9 binding sites. Analysis of the position and frequency of discrete rG4 forming sequences (*e.g*. G2N1, G2N3, or G2N5; Figure 5c and **Extended Figure 10b-c**) revealed a consistent enrichment of these motifs ~ 40 nt upstream of DHX9 peaks centre. Alignment of these motifs revealed G4-consensus motifs with defined G-tracts and short connecting loops (Figure 5d and **Extended Figure 10d-e**) suggesting that DHX9 binds downstream rG4 motifs. This result was further supported by the enrichment of sequences with lower than background predicted rG4 folding energies, *i.e*. predicted stable rG4 motifs, in the vicinity of all identified DHX9 peaks (**Extended Figure 10f**) and upstream DHX9 peaks within 5’-UTR (Figure 5e). It is noteworthy that biochemical analyses of DEAH-box helicases have shown that efficient loading to rG4 substrates require 15 nucleotides downstream the rG4 structural motif and that the helicases translocate in the 3’ to 5’ direction.^38,39^ These observations are consistent with our finding that DHX9 binds 40 nt downstream of ~ 25 nt long rG4 motifs in a cellular environment.

In order to link DHX9 binding to the change of TE upon DHX9 depletion, we analysed the ribosome distribution of transcripts displaying DHX9 iCLIP peaks within their 5’-UTR. Figure 5f **and 5g** report the DHX9 iCLIP signal together with ribosome occupancy along the *DDX23* and *COX14* transcripts. Both transcripts showed RPF enrichment upstream of rG4-containing DHX9 binding sites within their 5’-UTR. Upon depletion of DHX9, ribosome occupancy within 5’-UTRs increased while it decreased within the associated CDSs. Overall, transcripts bound by DHX9 were likewise characterised by negative and positive fold changes of TE and RPFdist respectively (Figure 5h-i). We also found an enrichment of the TE_down_ – RPFdist_up_ top motif upstream of 5’-UTR DHX9 peaks (**Extended Figure 10g-h**). It is noteworthy that transcripts bound by DHX9 in their 3’-UTR displayed DHX9-dependent translation (Figure 5h-i), which suggests a role of DHX9/rG4 in coordinating eIF4E and PABP1 during translation initiation.^40^ Taken together these observations demonstrate a direct regulation of ribosome distribution and TE by DHX9 through direct binding to its rG4 substrate.

### DHX36- and DHX9-dependent transcripts

Gene ontology classification for TE_down_ genes in DHX36 and DXH9 depleted samples (Figure 6a) revealed a preponderance of factors involved in gene expression regulation, chromatin remodelling and DNA damage/repair. Furthermore, we noted a significant enrichment of proto-oncogenes, such as MDM2, EGFR or CCAR2. Ranking of genes dependent on both helicases highlighted a consistent enrichment of transcription factors (*e.g*. STAT6 or FOXM1), epigenetic regulators (*e.g*. SUZ12, MLL1 or MLL5) and kinases (*e.g*. MAPK3, MAP2K1 or CDC42BPB) in both TE_down_ and RPFdist_up_ groups (Figure 6b-c). The individual RPF density plots illustrate recurrent patterns of altered ribosome distribution, whereas house-keeping genes (β-Actin, GAPDH and α-Tubulin) show no changes in ribosome distribution profile (**Extended Figure 11a**). We confirmed the impact of DHX36 and DHX9 depletion on key target proteins (Figure 6d and **Extended Figure 11b**), whilst controlling that the corresponding mRNAs were unaffected (**Extended Figure 11c**).

**Figure 6.**
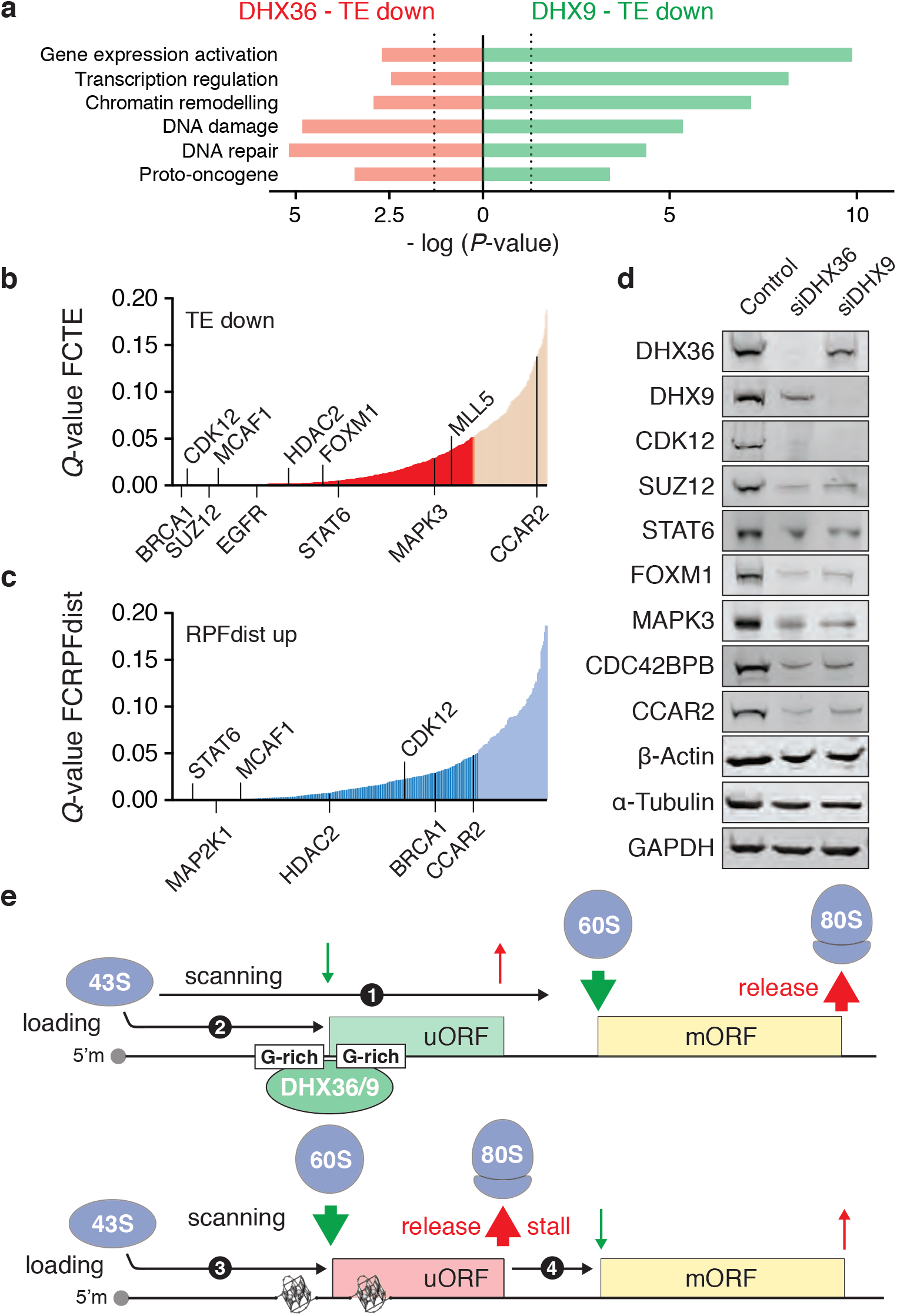
I DHX36 and DHX9 mediate translation of selected cancer genes. **a)** Gene ontology classification for genes, which TE decreases (*P* < 0.05) upon depletion of DHX36 (red) and DHX9 (green). **b)** TE_down_ (Q < 0.05) and (**c**) RPFdist_up_ (*Q* < 0.05) genes ranked by significance. TE_down_ and RPFdist_up_ transcripts include many genes involved in cancer pathways. **d)** Immunoblots of lysates from HeLa cells depleted in DHX36 and DHX9 and probed as indicated. Immunoblots were performed 96 h after siRNA transfection. **e)** Diagram showing a DHX36 / DHX9-dependent mechanism of translational control. 1. Scanning 43S PICs that translate unstructured 5’-UTRs or rG4-containning 5’-UTRs, that are maintained in their unfolded state by the DHX36 and DHX9 helicases, initiate translation at the main ORF (mORF). 2. A fraction of scanning PICs may initiate translation at upstream start codons, present within 5’-UTR in a suboptimal context, affecting the efficiency of the mORF translation. 3. In the absence of the rG4 processing helicases, rG4 motifs folding may slow down PIC scanning, thereby providing more time for the recognition of the upstream start codon and stimulating the translation of the upstream open reading frame (uORF). 4. 80S ribosomes may either dissociate from the mRNA after termination or stall during elongation or termination by the uORF-encoded attenuator peptide, preventing the translation of the mORF.

Because the DHX36- and DHX9-dependent transcripts included many genes with known role in cancer pathways, such as MAPK3/ERK1 or FOXM1,^41,42^ we sought to assess the mutation and expression profiles of DHX36 and DHX9 in cancer tissues to evaluate a potential contribution of both helicases in the oncogenic process. While we did not find any recurrent nor frequent mutations associated with DHX36 and DHX9 in cancer tissues (**Extended Figure 12a**), we found that human cancers show altered expression levels of both helicases. When comparing the expression levels of the helicases across normal tissue and tumours, recovered from the GENT database,^43^ we found that the helicases displayed altered expression levels in 8 and 9 cancers over the 15 types of cancer analysed for DHX36 and DHX9 respectively (**Extended Figure 12b-c**). When dysregulated, both helicases showed higher expression levels in tumours than in normal tissue (**Extended Figure 12d-e**) suggesting a role of both helicases in stimulating cancer pathways.

## DISCUSSION

Post-transcriptional regulation of gene expression allows a cell to orchestrate rapid changes in protein levels from steady state levels of mRNA. Cells have evolved *cis*-regulatory elements that are used to fine-tune the control of translation. Recent evidence has provided support that non-canonical secondary structures, such as rG4s, contribute to this mechanism by, for example, conferring eIF4A-dependent translation initiation^11^ or by impeding ribosome translocation.^44^ Our observations support an important role for 5’-UTR secondary structures in regulating translation initiation whilst revealing a particular effect of rG4 on 5’-UTR translation. We found that rG4 structures in mRNAs are associated with altered ribosome distribution and mark uORFs that display signs of active translation and thwart the translation of the downstream CDS. We propose that rG4 formation within 5’-UTRs impedes PIC scanning, thus promoting 80S ribosome formation upstream of canonical start codons (Figure 6e). This model is reminiscent of studies from M. Kozak showing that downstream secondary structures facilitate recognition of initiator codons by eukaryotic ribosomes.^45^ We tested this model, by studying the effect of two DEAH-box helicases, DHX36 and DHX9, on translation efficiency. Our experiments suggest that efficient processing of rG4 structures within 5’-UTR is required in order to favour translation at canonical start codons. It is noteworthy that the depletion of two RNA helicase paralogs affects the same subset of mRNAs, which demonstrate that mammalian cells have preserved their critical ability to process rG4 structures through evolution.

We found that rG4 structures can stimulate the translation of short open ORFs controlled by AUG, and non-AUG, codons occurring within 5’-UTRs. Independent of whether rG4-dependent uORFs are translated into stable peptides or N-terminal extensions, they could serve an important regulatory role. This is of special interest because of the growing evidence supporting a role of 5’-UTR translation on influencing human phenotypes and diseases. Indeed, besides the well-known role of uORFs in the integrated stress response pathway,^46^ polymorphic uORFs have been linked to gene expression variation^47^ and 5’-UTR translation to tumour initiation.^29^ The fact that DHX36 and DHX9 are dysregulated in cancer tissues supports the proposed role of RNA helicases in tumour initiation, progression and maintenance.^48^ This finding also supports that DHX36 and DHX9 should be included on the growing list of non-oncogenes that could be exploited as cancer drug targets. Owing to the functional redundancy of both helicases, they may not be essential in somatic cells for survival, but could become crucial in tumours in the absence of other rG4 processing factors establishing a “non-oncogene addiction” state.^49^

Characterising how two rG4-associated helicases modulate translation efficiency also supports that the rG4 structure rather than its nucleotide sequence repress translation. It has been recently proposed that rG4s are globally unfolded in eukaryotic cells, and that the regulatory role of rG4s in 5’-UTRs might result from the stable association of RNA-binding proteins that maintain the rG4 regions in the unfolded state.^50^ Our work shows that depletion of rG4 processing enzymes, that bind the structural motif in a cellular environment, causes changes in mRNA translation that are associated with rG4 structures demonstrating that rG4 folding can affect PIC scanning. Thus factors that modulate the folding/unfolding of rG4s can profoundly affect the translational landscape in human cells.

## Author contribution

PM designed the project and wrote the manuscript with contributions from all authors. PM performed the ribosome profiling and iCLIP experiments. PM, GM and AG analysed ribosome profiling and iCLIP datasets. PM and BH perfomed the polysome profiling coupled to mass spectrometry and affinity purification experiments. PM and GP performed statistical modelling. All authors contributed to analysing the data. SB supervised the project.

## Data accessibility

Ribosome profiling, matched mRNA-seq and iCLIP sequencing data have been deposited in the Gene Expression Omnibus database under accession number GSE105082.

## Acknowledgments

SB is a Wellcome Trust Senior Investigator (grant no. 099232/z/12/z). The Balasubramanian group is supported by a European Research Council Advanced Grant (no. 339778) and receives core funding (C14303/A17197) and programme funding (C9681/A18618) from Cancer Research UK.

## MATERIAL AND METHODS

### Ribosome profiling

HeLa cells were cultured in Dulbecco’s modified Eagle Medium (Life Technology) supplemented with 10% FBS (Life Technology). For siRNA transfection, 5 × 10^6^ HeLa cells per 15 cm dish were cultured overnight and treated with 27 μL DharmaFECT 1 Transfection Reagent (Dharmacon) and treated with 90 nM indicated siRNA in 1.8 mL OptiMEM media. Previously described DHX36 siRNA^1,2^ (GGGAACUGCGAA GAAGGUAUU, GUAAGGGAACUGCGAAGAA and CGGCAUGUGGUACGC GAAA) and DHX9 siRNA (UAGAAUGGGUGGAGAAGAAUU and GGCUAUAUCCAUCGAAAUUUU) were transfected at equimolar concentrations. A pool of four non-targeting siRNAs (ON-TARGETplus Nontargeting Control Pool, Dharmacon) was used as control. The cells were then expended for 48 h. DHX36 and DHX9 down-regulation was verified by immunoblot (see corresponding section). The cells were treated with cycloheximide (final concentration 0.1 mg.mL^−1^) for 1 min prior lysis.

Total RNA and ribosome-protected fragments (RPFs) were isolated using the TruSeq Ribo Profile Mammalian Kit (Illumina) following the manufacturer’s intructions with some modifications. For RPFs purification, lysates were treated with 5 U TruSeq Ribo Profile Nuclease per A260 of lysate. RPFs were isolated using Illustra MicroSpin S-400 HR Columns (GE Healthcare, 27-5140-01). rRNAs were removed using the Ribo-zero Magnetic Gold kit (Illumina, MRZG12324). After PAGE purification (15% urea-polyacrylamide gel to select fragments ~28–30 nt in length), end-repair, 3’-adapter ligation, reverse transcription (EpiScript reverse transcriptase) and cDNA circularisation (CircLigase, Epicentre), RPFs were amplified using 9 PCR cycles using Phusion polymerase (NEB). For parallel RNA-seq, total RNA was processed similarly, but excluding the nuclease digestion and S-400 columns purification steps. Before the library preparation steps, total RNA samples were heat-fragmented (94°C for 25 minutes). PCR amplicons were purified on a 8% native polyacrylamide gel excising bands corresponding to ~70–80 nt and ~80–100 nt for RPF and total RNA samples respectively. Libraries were sequenced on a NextSeq 500 using 75-nt single-read sequencing runs.

### Sequencing alignment and mapping

RNA-seq libraries were trimmed using cutadapt.^3^ The remaining reads were aligned to the GRCh38 human genome using the rsem-calculate-expression of RSEM^4^. For all analyses we used the version 26 of the human genome transcript annotation from Gencode. For each experiment, the transcriptome was defined by considering transcripts with an expression above 1 TPM (Transcript Per Million) and isoform representation percentage above 5%. This yielded the following number of selected transcripts for each different experiment: 11,557 for untreated cells; 18,455 for control siRNA-treated cells; 18,115 for siDHX36-treated cells and 20,439 for siDHX9-treated cells. 22,911 transcripts representing the union transcriptome of all performed RNA-seq experiments were then considered for all following analyses, and the corresponding RPF libraries were then aligned to this reference transcriptome using RSEM. Only RPF reads in the size range 24-32 were considered for analysis. Isoforms quantifications for both RNA-seq and RPF libraries were obtained in forms of estimated counts and TPM during the alignment procedure. Transcript coverage was calculated using RSEM command rsem-bam2wig and rsem-bam2readdepth, which both take into account multi-mapping reads and proportionally assign multiple assignments, a crucial step for processing short reads libraries.^4^ The coverage plots were then used to calculate total signal at 5’-UTR, CDS and 3’-UTR. Translation efficiency (TE) was calculated as the ratio of the total RPF signal (in TPM) over the coding region (CDS) divided by the total RNA signal (in TPM) over the same region: TE = RPF(CDS) / RNA(CDS). The 5’-UTR pattern of ribosomal occupancy, *i. e*. ribosome distribution, was defined as the ratio of the total RPF signal (in counts) over the 5’-UTR divided by the total RPF signal (in counts) over the associated CDS: RPFdist = RPF(5’-UTR) / RPF(CDS). Detailed results have been included in **Supplementary File 1**.

### Differential translation and ribosome distribution analysis

Change in translation efficiency upon depletion of the DHX9 and DHX36 helicases was defined as TE_siDHX36_/TE_control_ and TE_siDHx9_/TE_control_ respectively, where TE_control_ are values obtained from control siRNA-treated cells. To assess statistical relevance of changes in translation efficiency, RPF and total RNA signals (in TPM) were taken into Xtail,^5^ an *R* package designed for the identification of differentially translated genes in pairwise comparisons. Xtail estimates and accounts for biological variability in a statistical test based on the negative binomial distribution of the log2 fold changes in RPF and RNA signals. In our hands, Xtail was found to exhibit higher sensitivity as compared to other available pipelines, such as DESeq2^6^ or RiboDiff^7^. To assess co-change in translation efficiency upon depletion of both helicases, *P*-values (assessed using Xtail) were combined using Fisher’s method, and referred to as *Q*-values.^8^ Change in ribosome distribution was defined as RPFdistsiDHX36 / RPFdistcontrol and RPFdistsiDHX9 / RPFdistcontrol, where RPFdistcontrol are values obtained from control siRNA-treated cells. 5’-UTR and CDS RPF signal (in counts) were taken to DESeq2, as Xtail statistical model is tailored to translation efficiency estimation. *P*-values and log_2_ fold changes estimated by each method were used for all the following analyses. Detailed results have been included in Supplementary File 2. The TE_down_ – RPFdist_up_ and TE_up_ – RPFdist_down_ groups were defined by selecting transcripts with decreased or increased TE (*Q*-values ≤ 0.05) and increased or decreased RPFdist respectively.

### Meta-transcript profiles

Coverage files from estimated counts, generated by the alignment software RSEM (rsem-bam2readdepth command), were considered for the following analysis. To generate normalized profiles, total estimated counts, used for coverage normalization, were calculated for each library by summing up all transcript-wise estimated counts and dividing by 10^6^. The coverage profiles were calculated independently for 5’-UTR, CDS and 3’-UTR by sampling the normalized profile signal in 15, 90 and 75 bins, respectively. The bin numbers were chosen to reflect the average length distribution of the corresponding regions in all identified HeLa transcripts (22,911 transcripts representing the union transcriptome). Normalized transcripts were averaged together in a vectored way to plot the coverage distribution. Total RNA and RPF signals were treated separately in the same way. To generate coverage plots, outlier values, exceeding the 99.9 percentile for total RNA and RPF signals, were removed and the signal was normalised by the area under the curve. For condition-averaged profiles, the profiles of individual libraries were first averaged and then processed similarly.

### Predictors of translation initiation and efficiency

5’-UTR sequences using annotation from the version 26 of the human transcriptome from Gencode were recovered. These sequences were used to calculate the quantitative parameters used to describe the different mRNA features discussed in this manuscript. A complete and comprehensive list of these features is reported in the **Supplementary Methods**. RNA secondary structures were predicted using the RNAfold 2.2.10 algorithm of the ViennaRNA package.^9^ RNAfold computes the minimum free energy (MFE) of optimal secondary structures base on estimating base pairing probabilities. MFEs of dsRNA secondary structures (ΔG^0^_dsRNA_) were computed at 37°C. MFEs of rG4 secondary structures (ΔG^0^_rG4_) were computed by subtracting MFEs obtained when considering rG4 formation into the structure prediction algorithm to the previous values (ΔG^0^_rG4_ = ΔG^0^_dsRNA_ – ΔG^0^_dsRNA + rG4_).

### Hierarchical clustering analysis

To find groups of similar transcripts or similar helicases, hierarchical clustering was performed using the *R* environment. For transcripts clustering, log_2_ TE, 5’-UTR length normalised ΔG^0^_dsRNA_ and ΔG^0^_rG4_ values were used as z-scores. Different approaches were used to select the best clustering algorithm and to choose the optimal number of clusters. The reported analysis is using ‘canberra’ distance metrics and ‘ward.D2’ cluster method. Clusters were defined by identifying the five main groups of ΔG^0^_rG4_ values associated with different TE and ΔG^0^_dsRNA_. For helicases clustering, enrichments of each helicases in the three fractions (supernatant, monosomes, polysomes) were used as relative fractions (see **NanoLC–MS/MS analysis of Polysomal fractions** section for calculation details). The reported analysis used ‘euclidean’ distance metrics and ‘ward.D2’ cluster method.

### Principal Component Analysis and Statistical Modelling

Principal component analysis and the selection of predictive models were performed using the ‘factoextra’ ^10^ and ‘caret’ ^11^ package respectively in the *R* environment. A detailed and comprehensive description of the used pipeline is reported in the **Supplementary Methods**.

### Polysome profiling

Polysome analysis was performed as previously described.^12^ HeLa cells were grown in 15 cm dishes to 80% confluency. Cells were washed three times in cold PBS containing 100 μg.mL^−1^ of cycloheximide and scraped off the plate using a rubber policeman and 1mL of the same solution. Cells were centrifuged for 5 min at 1,000 rpm and resuspended in 425 μL of hypotonic lysis buffer (5 mM Tris-HCl pH 7.5, 2.5 mM MgCl_2_, 1.5 mM KCl) supplemented with 25 μL 10% TritonX100, 25 μL 10% Sodiumdeoxicholate, 1 μL 1M DTT and 5 μL RNAase inhibitor (40 U/μL, Promega). The supernatant was loaded onto a 10%-50% sucrose gradient prepared in 20 mM HEPES-KOH pH 7.6, 100 mM KCl and 5mM MgCl_2_ and was centrifuged in an SW40 rotor at 35,000 rpm for 2h. Gradients were analysed by piercing the tube with a Brandel tube piercer, passing 60% sucrose through the bottom of the tube, and monitoring the absorbance of the eluting material with an ISCO UA-6 UV detector. The different collected fractions were either analysed by mass spectrometry or by immunoblotting.

### NanoLC–MS/MS analysis of Polysomal fractions

Two biological replicates were performed. 30 μL of each polysome profiling fraction were loaded onto a 4-12% NuPAGE Bis-Tris gel that was run for 60 min at 180V and stained with Coomassie Brilliant Blue. Band corresponding to proteins and protein complexes with molecular weight above 100 kDa were excised (see **Extended** Figure 5a) and analysed by mass spectrometry as previously described.^13^ Briefly, protein bands were excised, and following several washes, the gel pieces were subjected to a reduction step using 10 mM DTT in 100 mM ammonium bicarbonate (NH4HCO3) buffer for 45 min at 56°C. Alkylation was performed using 55 mM iodoacetamide in 100 mM NH_4_HCO_3_ buffer for 30 min at room temperature in the dark. Digestion was performed using 10 μL of trypsin (10 mg/L in 50 mM NH_4_HCO_3_ buffer) overnight at 37°C. Eluted peptides were recovered, and the gel pieces were subsequently washed in 2.5% formic acid in 80% aqueous acetonitrile for 30 min at 37°C. The acid wash was combined with the original peptide eluate and dried. Samples were resuspended in 0.1% formic acid and analysed directly by nano-LC-MS/MS.

Mass spectrometry (MS) was performed using an LTQ Velos-Orbitrap MS (Thermo Scientific) coupled to an Ultimate RSLCnano-LC system (Dionex). Optimal separation conditions resulting in maximal peptide coverage was achieved using an Acclaim PepMap 100 column (C18, 3 μm, 100 Å) (Dionex) with an internal diameter of 75 μm and capillary length of 25 cm. A flow rate of 300 nL/min was used with a solvent gradient of 5% B to 40% B in 55 min followed by increasing the gradient to 95% B over 10 min. Solvent A was 0.1% (*v/v*) formic acid, 5% DMSO in water, whereas the composition of solvent B was 80% (v/v) acetonitrile, 0.1% (v/v) formic acid, 5% DMSO in water. The mass spectrometer was operated in positive ion mode using a Nth order double-play method to automatically switch between Orbitrap-MS and LTQ Velos-MS/MS acquisition. Survey full-scan MS spectra (from 400 to 1,600 m/z) were acquired in the Orbitrap with resolution (R) 60,000 at 400 m/z (after accumulation to a target of 1,000,000 charges in the LTQ). The method used allowed sequential isolation of the 20 most intense ions for fragmentation in the linear ion trap, depending on signal intensity, using CID at a target value of 3,000 charges. For accurate mass measurements, the lock mass option was enabled in MS mode, and the 445.120025 ion was used for internal recalibration during the analysis. Target ions already selected for MS/MS were dynamically excluded for 30 s. General MS conditions were electrospray voltage, 1.50 kV with no sheath or auxiliary gas flow, an ion selection threshold of 1,000 counts for MS/MS, an activation Q value of 0.25, activation time of 12 ms, capillary temperature of 200 °C, and an S-Lens RF level of 60 % were also applied. Charge state screening was enabled, and precursors with unknown charge state or a charge state of 1 were excluded. Raw MS data files were processed using Proteome Discoverer v.1.4 (Thermo Scientific). Processed files were searched against the SwissProt human database using the Mascot search engine version 2.3.0. Searches were done with tryptic specificity allowing up to one miscleavage and a tolerance on mass measurement of 10 ppm in MS mode and 0.6 Da for MS/MS ions. Structure modifications allowed were oxidized methionine, and deamidation of asparagine and glutamine residues, which were searched as variable modifications. Using a reversed decoy database, false discovery rate (FDR) was less than 1%. Detailed results have been included in **Supplementary File 3**.

The presence of proteins in supernatant, monosome and polysome fractions was qualitatively assessed by analysing the number of unique peptides in each sample. Functional analysis of proteins was performed using the DAVID bioinformatics resources 6.8^14^ using all human proteins with molecular weight above 100 kDa as background. Quantitative enrichment of helicases in polysome fractions was assessed by calculating the ratio of number of unique peptides in each fraction (supernatant, monosome or polysome) over the total number of unique peptides in all fractions. To avoid false positive, proteins with less than 10 unique peptides in all fractions were excluded from the analysis. Calculated relative fractions were used for the clustering analysis (see **Hierarchical clustering analysis section**).

### Immunoblots and Antibodies

Effects of siRNA transfection on DHX36 and DHX9 protein levels were assessed after 48h. Down-stream effect of DHX36 and DHX9 depletion was tested after 96h and two consecutive rounds of siRNA transfections. siRNA transfection was performed as described previously (see **Ribosome profiling** section). Total cell lysates were prepared using Laemmli lysis buffer (92 mM Tris.HCl pH 6.8, 18 % glycerol, 1.8 % SDS, 0.02 % Bromophenol Blue, 2 % β-mercaptoethanol). Before lysis, cells were harvested by trypsinization, suspended in media with serum and counted. After centrifugation (5 min at 1,000 rpm), the cell pellets were resuspended in Laemmli lysis buffer at a concentration of 10^7^ cells per mL, heated at 95°C for 5 minutes and sonicated. The equivalent of 10^5^ cells (10 μL) was loaded onto 4-12% NuPAGE Bis-Tris gels then transferred onto nitro-cellulose membranes using an iBlot 2 Gel Transfer Device (ThermoFisher Scientific). Membranes were blocked for 60 min in Odyssey Blocking Buffer (LI-COR Biosciences) and incubated overnight at 4°C in solutions of primaries antibodies in the blocking buffer. After washing with TBS supplemented with 0.1% Tween 20, membranes were incubated with secondary antibodies in the blocking buffer at room temperature for 60 min. IRDye secondary antibodies (LI-COR Biosciences) were used to detect protein bands on a Odyssey CLx Imaging System (LI-COR Biosciences).

Primary antibodies used in this study were DHX36 (Abcam ab70269), DHX9 (RNA Helicase A, Abcam ab26271), DHX29 (Abcam ab70745), DHX30 (Abcam ab85687), DHX57 (Abcam ab86784), TDRD9 (Abcam ab118427), YTHDC2 (Abcam ab176846), S6 ribosomal protein (Cell Signaling #2217), CDK12 (CRKRS, Abcam ab57311), SUZ12 (Abcam ab175187), STAT6 (Abcam ab32520), FOXM1 (Santa Cruz Biotechnology sc-376471), MAPK3 (ERK1, Abcam ab32537), CDC42BPB (Abcam ab61328), CCAR2 (KIAA1967, Abcam ab205526), β-Actin (ACTB, Cell Signalling #4970), α-Tubulin (TUBA1A, Cell Signalling #86298), GAPDH (Sigma-Aldrich G8795).

### Affinity purification

Affinity purifications were performed as previously described with the following adjustments.^15^ HeLa cells (4 × 10^6^ cells) were lysed in hypotonic lysis buffer as described and the protein concentration was determined with the BioRad protein assay (BiorRad) according to the manufacturer’s suggestions. One mg of cytoplasmic extracts was precleared with Streptavidin MagneSphere^®^ Paramagnetic Particles (Promega) in the RNA pull down buffer (20 mM Hepes pH8, 100 mM NaCl, 20% glycerol, 0.2 mM EDTA, 1 mM DTT, 0.01 % Nonidet-P40, 50 μg/mL yeast tRNA (Ambion), 160 U/mL RNasin). Prior to enrichements, 10 μM solutions of biotinylated oligonucletides were annealed in 1x PBS supplemented with 2M KCl by boiling for 5 minutes and slowly cooling to room temperature and kept at 4 °C until use. The nucleic acids probes were, rG4: (Biotin-UGUGGGAGGGGCGGGUCUGGGUGC), mG4 (Biotin-UGUAGAAAGAGCAGAUCUAGAUGC) and SL (Biotin-ACAGGGCUCCGCGAUGGCGGAGCCCAA). Empty beads (B) were used as negative control. Biotinylated RNAs (5nM) were bound to streptavidin beads and afterwards combined with precleared cytoplasmic extracts to perform affinity purifications for 4 hours at 4 °C. The beads were washed with RNA pull down buffer three times and one time with 1x PBS. Interacting proteins were eluted by boiling in 50 μL 1x SDS Laemmli buffer. One half of the eluted protein complexes in Laemmli buffer were loaded onto a 4-12% NuPAGE Bis-Tris gel and the helicases of interest interrogated by immunoblotting with specific antibodies (see **Immunoblots and Antibodies section**).

Motif identification and analysis. De novo motif discovery and analysis were performed using the Meme suite.^16^ The 5’-UTR sequences of the TE_down_ – RPFdist_up_ and TE_up_ – RPFdist_down_ sets were collected for motifs prediction. The Meme tool was run using both sequence datasets as primary sequences and the 5’-UTR sequences of all HeLa transcripts, as assessed by RNA-seq, were used as control sequences. A strand-specific 3-order Markov model was used to correct for biased frequencies of all k-mers (k ≤ 3). Meme was run to identify enriched motifs of a maximum length of 30 nucleotides that occur any number of time in a given 5’-UTR sequence. The occurrence of the five more enriched motifs in the primary set of sequences, as compared to the control set, were called using FIMO with default parameters for strand specific prediction correcting for biased frequencies of all k-mers (k ≤ 3). Enrichment of motifs was calculated by comparing the density of the motifs to the density of the same motif in the unchanged set, defined as transcripts with FC TE between −0.1 and 0.1. Density of motifs is defined as the ratio of the number of motif occurrence and the total number of residues in a given set. *P*-values for motif enrichment were calculated using a two-sided Fisher exact test. For base composition analysis and predicted secondary structure prediction, the sequences of the most enriched motif in the TE_down_ – RPFdist_up_ set were recovered together with a 10 nucleotides flanking region. *P*-values for difference in base composition and stability of predicted structures were assessed using a Mann–Whitney test.

### Identification of uORFs and ORFscore calculation

Translated 5’-leader sequences in ribosome profiling data from HeLa and DHX36/DHX9-depleted HeLa cells were predicted using the ORFscore pipeline.^17^ uORFs were defined as sequences in annotated 5’-UTRs (Gencode version 26) with start codons (AUG, CUG, UUG and GUG) in frame with a stop codon (UAA, UAG and UGA). An ORFscore was then calculated for each identified uORFs. To calculate the ORFscores, 28-29 nt RPF reads of each replicates were combined and counted at each position within the uORFs, excluding the first and last coding codons. Any uORFs without RPF reads in one of the replicate were excluded from the analysis. To avoid false positives due to little information about RPF positioning, uORFs with low read coverage (< 350 reads per kilobases) were excluded from the analysis. The ORFscore was then calculated as:

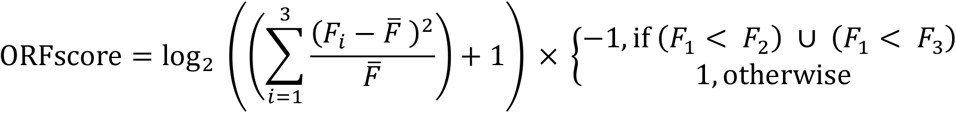

where *F_n_* is the number of reads in reading frame *n*, 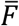 is the total number of reads across all three frames divided by 3. Hence the RPF distribution in each frame of a given uORF is compared to an equally sized uniform distribution using a modified chi-squared statistic. Negative and positive ORFscores are assigned when the distribution of RPFs is inconsistent or consistent, respectively, with the frame of a given uORF. An ORFscore threshold of 6 was used to call with confidence for uORFs that are translated since annotated CDS regions are characterised by higher than this threshold ORFscores.^17^ Detailed results have been included in **Supplementary File 4**.

### iCLIP

Two biological replicates of DHX9 iCLIP were performed as previously described,^18^ with some modifications. For each experiments, 8 150mm plates of HeLa cells were seeded to be at ~90% confluence during UV crosslinking. For crosslinking, the cells were washed with cold PBS and then the plates where irradiated on ice with 150 mJ/cm2 at 254nm. The cells were scraped into PBS, pelleted by centrifugation at 1,000g for 5min at 4°C, and the pellets were resuspended in 500 μL lysis buffer (50mM Tris-HCl, pH 7.4, 100 mM NaCl, 1% Igepal CA-360, 0.1% SDS, 0.5% sodium deoxycholate, 1/100 volume Protease Inhibitor Cocktail). At this stage, 4 lysates were combined to generate 4 samples for RNA digestion and immunoprecipitation. RNA digestion was performed using 4 unit of RNAse I (Life Technologies AM2295) per mL of lysate. Immunoprecipitation was perform using 100 μL of protein A-coated Dynabeads (Life Technologies 10002D) and a DHX9 antibody (Abcam ab26271) per IP. Beads were extensively washed with high-salt buffer (50mM Tris-HCl, pH 7.4, 1 M NaCl, 1 mM EDTA, 1% Igepal CA-360, 0.1% SDS, 0.5% sodium deoxycholate). After 3’-end dephosphorylation, 3’ adaptor ligation and ^32^P 5’-end labelling, a small aliquot (10% of the total volume) was saved for immunoblot analysis while the remaining samples were loaded onto a 4-12% NuPAGE Bis-Tris gel that was run for 60 min at 180V. The RNA-protein complexes were transfered to a Protan BA85 Nitrocellulose Membrane using a Novex wet transfer apparatus for 2 h at 30 V. After transfer, the membrane was rinsed in PBS buffer and exposed to a Fuji film at −20°C overnight. Immunoblots were used to identify the DHX9-RNA complexes to be isolated from the membrane. At this stage two nitrocellulose pieces were combined to generate the two biological replicates. Each samples were then incubated for 60 min at 50°C in 200 μL PK/SDS buffer (100 mM Tris, pH 7.5, 50 mM NaCl, 1 mM EDTA, 0.2% SDS) supplemented with 10 μL proteinase K (Fisher Scientific YSJ-762-Q). RNA-protein complexes were recovered by Phenol:Chloroform extraction and ethanol precipitation. Reverse transcription was performed using Superscript III (Life Technologies 18080085) following the manufacturer’s instructions. Residual RNA was removed by alkaline hydrolysis. cDNA was size selected on a 6% polyacrylamide / 7-8 M Urea / TBE gel run at 180 V for 40 min. Bands equivalent of 80-120 nt were excised from the gel. PCR amplification was performed using Accuprime Supermix I (Life Technologies 12342028) and 20 to 24 cycles to avoid the formation of secondary products. PCR products were purified on a 8% native polyacrylamide gel run at 200V for 30 min. Amplicons of 140-170nt were excised from the gel and recover by ethanol precipitation. PCR primers contained the Illumina P5 and P3 sequences together with degenerated barcodes for PCR duplicates removal. The iCLIP libraries were sequenced on a NextSeq 500 using 75-nt single-read sequencing runs.

Raw Illumina reads were processed as follow: Barcodes (NNNCGGANNN and NNNGGCANNN) were used for demultiplexing. PCR duplicates, *i.e*. reads having the same sequence and barcode, were removed using a customized Unix script. Remaining reads were aligned to the HeLa transcriptome (22,911 transcripts identified from RNA-seq experiments) using RSEM (rsem-calculate-expression command). Aligned reads were further processed for duplicates removal, *i.e*. reads aligning at the same location and having the same 10-mer barcode, leaving a single read per location. Transcript coverage was calculated using the rsem-bam2wig command, which takes into account and proportionally assigns multi-mapping reads. Bedgraph files were calculated (command bigWigToBedGraph from the UCSC utilities) and analysed for peak calling using MACS2^19^ (command macs2 bdgpeakcall -l 40 -g 30 -c 5). Multi-modal peaks were redefined after combining reads from both duplicates and were splitted using PeakSplitter,^20^ the middle points from each peak were extracted for further analysis. Detailed results have been included in **Supplementary File 5**.

### Statistics

Data were analysed and statistics performed in Prism6 (GraphPad) and the *R* environment. Significant differences between two groups were noted by asterisks (\**P* < 0.05, \*\**P* < 0.01, \*\*\**P* < 0.001). Replicates (n) in this study refer to biological replicates.

## REFERENCES

1. Weinberg, D. E., Shah, P., Eichhorn, S. W., Hussmann, J. A., Plotkin, J. B. & Bartel, D. P. Improved Ribosome-Footprint and mRNA Measurements Provide Insights into Dynamics and Regulation of Yeast Translation. Cell Rep. 14, 1787–1799 (2016).

2. Ingolia, N. T., Lareau, L. F. & Weissman, J. S. Ribosome profiling of mouse embryonic stem cells reveals the complexity and dynamics of mammalian proteomes. Cell 147, 789–802 (2011).

3. Truitt, M. L. & Ruggero, D. New frontiers in translational control of the cancer genome. Nat. Rev. Cancer 16, 288–304 (2016).

4. Hinnebusch, A. G., Ivanov, I. P. & Sonenberg, N. Translational control by 5’-untranslated regions of eukaryotic mRNAs. Science 352, 1413–1416 (2016).

5. Hinnebusch, A. G. Molecular Mechanism of Scanning and Start Codon Selection in Eukaryotes. Microbiol. Mol. Biol. Rev. 75, 434–467 (2011).

6. Dever, T. E. & Green, R. The Elongation, Termination, and Recycling Phases of Translation in Eukaryotes. Cold Spring Harb Perspect Biol 4:a013706 (2012).

7. Gingras, A., Raught, B. & Sonenberg, N. eIF4 Initiation Factors: Effectors of mRNA Recruitment to Ribosomes and Regulators of Translation. Annu. Rev. Biochem. 68, 913–963 (1999).

8. Chu, D. & Haar, T. Von Der. The architecture of eukaryotic translation. Nucleic Acids Res. 40, 10098–10106 (2017).

9. Shah, P., Ding, Y., Niemczyk, M., Kudla, G. & Plotkin, J. B. Rate-Limiting Steps in Yeast Protein Translation. Cell 153, 1589–1601 (2013).

10. Parsyan, A., Svitkin, Y., Shahbazian, D., Gkogkas, C., Lasko, P., Merrick, W. C. & Sonenberg, N. mRNA helicases: the tacticians of translational control. Nat. Rev. Mol. Cell Biol. 12, 235–45 (2011).

11. Wolfe, A. L. et al. RNA G-quadruplexes cause eIF4A-dependent oncogene translation in cancer. Nature 513, 65–70 (2014).

12. Rubio, C. A., Weisburd, B., Holderfield, M., Arias, C., Fang, E., DeRisi, J. L. & Fanidi, A. Transcriptome-wide characterization of the eIF4A signature highlights plasticity in translation regulation. Genome Biol. 15, 476 (2014).

13. Soto-Rifo, R., Rubilar, P. S., Limousin, T., de Breyne, S., Décimo, D. & Ohlmann, T. DEAD-box protein DDX3 associates with eIF4F to promote translation of selected mRNAs. EMBO J. 31, 3745–56 (2012).

14. Pisareva, V. P., Pisarev, A. V., Komar, A. a., Hellen, C. U. T. & Pestova, T. V. Translation Initiation on Mammalian mRNAs with Structured 5’UTRs Requires DExH-Box Protein DHX29. Cell 135, 1237–1250 19 (2008).

15. Hartman, T. R., Qian, S., Bolinger, C., Fernandez, S., Schoenberg, D. R. & Boris-Lawrie, K. RNA helicase A is necessary for translation of selected messenger RNAs. Nat. Struct. Mol. Biol. 13, 509–516 (2006).

16. Sen, N. D., Zhou, F., Ingolia, N. T. & Hinnebusch, A. G. Genome-wide analysis of translational efficiency reveals distinct but overlapping functions of yeast DEAD-box RNA helicases Ded1 and eIF4A. Genome Res. 25, 1196–1205 (2015).

17. Dabas, N., Zhou, F., Harris, M. S., Ingolia, N. T. & Hinnebusch, A. G. eIF4B stimulates translation of long mRNAs with structured 5 ′ UTRs and low closed-loop potential but weak dependence on eIF4G. Proc. Natl. Acad. Sci. 113, 10464–10472 (2016).

18. Mandel, J., Ehresmann, B. & Ehresmann, C. The fragile X mental retardation protein binds specifically to its mRNA via a purine quartet motif. EMBO J. 4813, 4803–4813 (2001).

19. Kumari, S., Bugaut, A., Huppert, J. L. & Balasubramanian, S. An RNA G-quadruplex in the 5′ UTR of the NRAS proto-oncogene modulates translation. Nat. Chem. Biol. 3, 218–221 (2007).

20. Bugaut, A. & Balasubramanian, S. 5’-UTR RNA G-quadruplexes: Translation regulation and targeting. Nucleic Acids Res. 40, 4727–4741 (2012).

21. Ingolia, N. T., Ghaemmaghami, S., Newman, J. R. S. & Weissman, J. S. Genome-Wide Analysis in Vivo of Translation with Nucleotide Resolution Using Ribosome Profiling. Science 324, 218–224 (2009).

22. Brar, G. A., Yassour, M., Friedman, N., Regev, A., Ingolia, N. T. & Weissman, J. S. High-Resolution View of the Yeast Meiotic Program Revealed by Ribosome Profiling. Science 335, 552–558 (2012).

23. Ruggero, D. Translational control in cancer etiology. Cold Spring Harb. Perspect. Biol. 5, 1–27 (2013).

24. Bedrat, A., Lacroix, L. & Mergny, J. Re-evaluation of G-quadruplex propensity with G4Hunter. Nucleic Acids Res. 44, 1746–1759 (2016).

25. Truitt, M. L., Conn, C. S., Shi, Z., Seo, Y., Barna, M., Truitt, M. L., Conn, C. S., Shi, Z., Pang, X., Tokuyasu, T. & Coady, A. M. Differential Requirements for eIF4E Dose in Normal Development and Cancer. Cell 162, 59–71 (2015).

26. Hsieh, A. C. et al. The translational landscape of mTOR signalling steers cancer initiation and metastasis. Nature 485, 55–61 (2012).

27. Zur, H. & Tuller, T. New Universal Rules of Eukaryotic Translation Initiation Fidelity. PLOS Comput. Biol. 9, e1003136 (2013).

28. Nikolaev, S. I. et al. Exome sequencing identifies recurrent somatic MAP2K1 and MAP2K2 mutations in melanoma. Nat. Publ. Gr. 44, 133–20 139 (2011).

29. Sendoel, A., Dunn, J. G., Rodriguez, E. H., Naik, S., Gomez, N. C., Hurwitz, B., Levorse, J., Dill, B. D., Schramek, D., Molina, H., Weissman, J. S. & Fuchs, E. Translation from unconventional 5′ start sites drives tumour initiation. Nature 541, 494–499 (2017).

30. Bazzini, A. A., Johnstone, T. G., Christiano, R., Mackowiak, S. D., Obermayer, B., Fleming, E. S., Vejnar, C. E., Lee, M. T., Rajewsky, N., Walther, T. C. & Giraldez, A. J. Identification of small ORFs in vertebrates using ribosome footprinting and evolutionary conservation. EMBO J. 33, 981–993 (2014).

31. Pauli, A., Schier, A. F. & Chew, G. Conservation of uORF repressiveness and sequence features in mouse, human and zebrafish. Nat. Commun. 7, 11663 (2016).

32. Umate, P., Tuteja, N. & Tuteja, R. Genome-wide comprehensive analysis of human helicases. Commun. Integr. Biol. 4, 1–20 (2011).

33. Chakraborty, P. & Grosse, F. Human DHX9 helicase preferentially unwinds RNA-containing displacement loops (R-loops) and Gquadruplexes. DNA Repair 10, 654–665 (2011).

34. Creacy, S. D., Routh, E. D., Iwamoto, F., Nagamine, Y., Akman, S. A. & Vaughn, J. P. G4 Resolvase 1 Binds Both DNA and RNA Tetramolecular Quadruplex with High Affinity and Is the Major Source of Tetramolecular Quadruplex G4-DNA and G4-RNA Resolving Activity in HeLa Cell Lysates. J. Biol. Chem. 283, 34626–34634 (2008).

35. Thandapani, P., Song, J., Gandin, V., Cai, Y., Rouleau, S. G., Garant, J.-M., Boisvert, F.-M., Yu, Z., Perreault, J.-P., Topisirovic, I. & Richard, S. Aven recognition of RNA G-quadruplexes regulates translation of the mixed lineage leukemia protooncogenes. Elife 4, e06234 (2015).

36. Bailey, T. L., Boden, M., Buske, F. A., Frith, M., Grant, C. E., Clementi, L., Ren, J., Li, W. W. & Noble, W. S. MEME SUITE: tools for motif discovery and searching. Nucleic Acids Res. 37, 202–208 (2017).

37. Huppertz, I., Attig, J., D’Ambrogio, A., Easton, L. E., Sibley, C. R., Sugimoto, Y., Tajnik, M., Konig, J. & Ule, J. iCLIP: Protein-RNA interactions at nucleotide resolution. Methods 65, 274–287 (2014).

38. Giri, B., Smaldino, P. J., Thys, R. G., Creacy, S. D., Routh, E. D., Hantgan, R. R., Lattmann, S., Nagamine, Y., Akman, S. A. & Vaughn, J. P. G4 Resolvase 1 tightly binds and unwinds unimolecular G4-DNA. Nucleic Acids Res. 39, 7161–7178 (2011).

39. Tippana, R., Hwang, H., Opresko, P. L., Bohr, V. A. & Myong, S. Singlemolecule imaging reveals a common mechanism shared by Gquadruplex– resolving helicases. Proc. Natl. Acad. Sci. 113, 8448–8453 (2016).

40. Kahvejian, A., Svitkin, Y. V, Sukarieh, R., Boutchou, M. M. & Sonenberg, N. Mammalian poly (A)-binding protein is a eukaryotic translation initiation factor, which acts via multiple mechanisms. Genes Dev. 19, 104–113 (2005).

41. Kim, E. K. & Choi, E. Biochimica et Biophysica Acta Pathological roles of MAPK signaling pathways in human diseases. Biochim. Biophys. Acta 1802, 396–405 (2010).

42. Raychaudhuri, P. & Park, H. J. FoxM1: A Master Regulator of Tumor Metastasis. Cancer Res. 71, 4329–4334 (2011).

43. Shin, G., Kang, T., Yang, S., Baek, S. & Jeong, Y. GENT: Gene Expression Database of Normal and Tumor Tissues. Cancer Inform. 10, 149–157 (2011).

44. Murat, P., Zhong, J., Lekieffre, L., Cowieson, N. P., Clancy, J. L., Preiss, T., Balasubramanian, S., Khanna, R. & Tellam, J. G-quadruplexes regulate Epstein-Barr virus-encoded nuclear antigen 1 mRNA translation. Nat. Chem. Biol. 10, 358–64 (2014).

45. Kozak, M. Downstream secondary structure facilitates recognition of initiator codons by eukaryotic ribosomes. Proc. Natl. Acad. Sci. U. S. A. 87, 8301–8305 (1990).

46. Young, S. K. & Wek, R. C. Upstream Open Reading Frames Differentially Regulate Gene-specific Translation in the Integrated Stress Response. J. Biol. Chem. 291, 16927–16935 (2016).

47. Calvo, S. E., Pagliarini, D. J. & Mootha, V. K. Upstream open reading frames cause widespread reduction of protein expression and are polymorphic among humans. Proc Natl Acad Sci U S A 106, 7507–7512 (2009).

48. Robert, F. & Pelletier, J. Perturbations of RNA helicases in cancer. Wiley Interdiscip. Rev. RNA 4, 333–349 (2013).

49. Solimini, N. L., Luo, J. & Elledge, S. J. Non-Oncogene Addiction and the Stress Phenotype of Cancer Cells. Cell 130, 986–988 (2007).

50. Guo, J. U. & Bartel, D. P. RNA G-quadruplexes are globally unfolded in eukaryotic cells and depleted in bacteria. Science 353, aaf5371–6 (2016).

## SUPPLEMENTARY REFERENCES

1. Tran, H., Schilling, M., Wirbelauer, C., Hess, D. & Nagamine, Y. Facilitation of mRNA Deadenylation and Decay by the Exosome-Bound, DExH Protein RHAU. Mol. Cell 13, 101–111 (2004).

2. Kim, T., Pazhoor, S., Bao, M., Zhang, Z., Hanabuchi, S., Facchinetti, V. & Bover, L. Aspartate-glutamate-alanine-histidine box motif (DEAH)/ RNA helicase A helicases sense microbial DNA in human plasmacytoid dendritic cells. Proc Natl Acad Sci U S A 107, 15181–15186 (2010).

3. Martin, M. Cutadapt removes adapter sequences from high-throughput sequencing reads. EMBnet.journal 17, 10–12 (2011).

4. Li, B. & Dewey, C. N. RSEM: accurate transcript quantification from RNA-Seq data with or without a reference genome. BMC Bioinformatics 12, 323 (2011).

5. Xiao, Z., Zou, Q., Liu, Y. & Yang, X. Genome-wide assessment of differential translations with ribosome profiling data. Nat. Commun. 7, 1–11 (2016).

6. Love, M. I., Huber, W. & Anders, S. Moderated estimation of fold change and dispersion for RNA-seq data with DESeq2. Genome Biol. 15, 550 (2014).

7. Zhong, Y., Karaletsos, T., Drewe, P., Sreedharan, V. T., Kuo, D., Singh, K. & Wendel, H. Gene expression RiboDiff: detecting changes of mRNA translation efficiency from ribosome footprints. Bioinformatics 33, 139–141 (2017).

8. Fisher, R. A. Statistical Methods for Research Workers. (1932).

9. Package, V., Lorenz, R., Bernhart, S. H., Höner, C., Tafer, H. & Flamm, C. ViennaRNA Package 2.0. Algorithms Mol. Biol. 6, 26 (2011).

10. Kassambara, A. & Mundt, F. factoextra: Extract and Visualize the Results of Multivariate Data Analyses. R Packag. version 1.0.4. https://CRAN.R-project.org/package=factoextra (2017).

11. Kuhn, M. Building Predictive Models in R Using the caret Package. J. Stat. Softw. 28, (2008).

12. Costa-mattioli, M., Gobert, D., Stern, E., Gamache, K., Colina, R., Cuello, C., Sossin, W., Kaufman, R., Pelletier, J., Rosenblum, K., Krnjevic, K., Lacaille, J.-C., Nader, K. & Sonenberg, N. eIF2 a Phosphorylation Bidirectionally Regulates the Switch from Short-to Long-Term Synaptic Plasticity and Memory. Cell 129, 195–206 (2007).

13. Mohammed, H. et al. Report Endogenous Purification Reveals GREB1 as a Key Estrogen Receptor Regulatory Factor. Cell Rep. 3, 342–349 (2013).

14. Huang, D. W., Sherman, B. T. & Lempicki, R. A. Bioinformatics enrichment tools: paths toward the comprehensive functional analysis of large gene lists. Nucleic Acids Res. 37, 1–13 (2009).

15. Herdy, B., Karonitsch, T., Vladimer, G. I., Tan, C. S. H., Stukalov, A., Trefzer, C., Bigenzahn, J. W., Theil, T., Holinka, J., Kiener, H. P., Colinge, J., Bennett, K. L. & Superti-Furga, G. The RNA-binding protein HuR/ELAVL1 regulates IFN-β mRNA abundance and the type I IFN response. Eur. J. Immunol. 45, 1500–1511 (2015).

16. Bailey, T. L., Boden, M., Buske, F. A., Frith, M., Grant, C. E., Clementi, L., Ren, J., Li, W. W. & Noble, W. S. MEME SUITE: tools for motif discovery and searching. Nucleic Acids Res. 37, 202–208 (2017).

17. Bazzini, A. A., Johnstone, T. G., Christiano, R., Mackowiak, S. D., Obermayer, B., Fleming, E. S., Vejnar, C. E., Lee, M. T., Rajewsky, N., Walther, T. C. & Giraldez, A. J. Identification of small ORFs in vertebrates using ribosome footprinting and evolutionary conservation. EMBO J. 33, 981–993 (2014).

18. Huppertz, I., Attig, J., D’Ambrogio, A., Easton, L. E., Sibley, C. R., Sugimoto, Y., Tajnik, M., Konig, J. & Ule, J. iCLIP: Protein-RNA interactions at nucleotide resolution. Methods 65, 274–287 (2014).

19. Zhang, Y., Liu, T., Meyer, C. A., Eeckhoute, J., Johnson, D. S., Bernstein, B. E., Nusbaum, C., Myers, R. M., Brown, M., Li, W. & Liu, X. S. Open Access Model-based Analysis of ChIP-Seq (MACS). R137 (2008). doi:10.1186/gb-2008-9-9-r137

20. Dvinge, H., Tammoja, K. & Bertone, P. PeakAnalyzer: Genome-wide annotation of chromatin binding and modification loci. BMC Genomics 415–427 (2010).

